# Media Matters: Phenol Red and Fetal Bovine Serum Estrogen in Traditional Cell Culture Media Influence Human Mesenchymal Stromal Cell (hMSC) Processes and Differentiation in a Sex-Biased Manner

**DOI:** 10.1101/2025.05.27.656455

**Authors:** John C Bradford, Jennifer L Robinson

## Abstract

Estrogens are global regulators of cellular signaling pathways, impacting fundamental processes and phenotypes that are essential for tissue remodeling and homeostasis. Traditional cell culture media contains estrogen-mimetic compounds, including phenol red and endogenous estrogen in fetal bovine serum (FBS). However, the potential of these compounds to bias *in vitro* studies, particularly when considering sex as a biological variable, remains unclear. This gap in understanding critically impacts the culture of human mesenchymal stromal cells (hMSCs), whose basic functions and differentiation potential, central to cell therapy and tissue engineering, are sensitive to perturbations in the culture conditions. Despite this, the effect of estrogens from cell culture media on male and female hMSCs is not currently considered in cell processing for clinical trials. As such, a baseline understanding of these estrogen-mimetic media influences on hMSCs is critical for clinical efficacy and adequate study design in research. To this end, we investigated the effects of phenol red and fetal bovine serum on the proliferation, metabolism, senescence, and differentiation capacity of male and female hMSCs. Phenol red, FBS, donor sex, and 17β-estradiol (E2) supplementation all had significant impacts on hMSC health and differentiation potential in culture. Notably, dosing with estrogen at the levels found in FBS did not recover most of the hMSC metrics tested. The only outcomes that were not significantly different based on donor sex were senescence and mRNA transcripts for RUNX2 and PPARG, transcriptional regulators for osteogenesis and adipogenesis. Overall, these findings reveal the sex-biased effects of estrogen and estrogen-mimetic compounds in traditional culture media, underscoring a current gap in considering sex as a biological variable in cell therapy and tissue engineering research and manufacturing.

## Introduction

Males and females have varying exposure levels to estrogens across the lifespan (1) and male and female tissues exhibit variable responses to similar concentrations of estrogens (2). While it is widely known that 17β-estradiol (E2), the main estrogen isoform from puberty to menopause, impacts reproductive tissues, the it also controls important processes in other tissue systems including nervous, musculoskeletal, and immune systems (3–5). In these tissues, E2 has multiple mechanisms of action, including canonical genomic action, non-genomic membrane-initiated steroid signaling (MISS), and DNA methylation and epigenetic regulation. Because of this, care must be taken when analyzing human data to consider both the recent and historical levels of E2 exposure to fully account for differences arising from E2 signaling. During pre-pubescence, females have higher levels of E2 than males due to higher aromatase activity in the tissues, an effect exacerbated during pubescence with increased production of E2 in the ovaries (2,6). During perimenopause and menopause, E2 levels become highly variable, and while they tend to decrease overall, they can spike roughly 3 times higher than their peak levels seen in pre-menopausal ovulatory cycles (7,8). Following menopause, females revert to their pre-pubescent methods of E2 production, predominantly occurring outside of the ovaries from aromatase conversion of testosterone (2). These varying levels affect both the nascent phenotype and the long-term epigenetic programming of hMSCs (9,10), motivating the need to characterize the impact of E2 for both primary research and clinical studies on cell therapy applications. Due to the historical exclusion of females from research, in 2016 the NIH implemented the sex as a biological variable (SABV) policy. While this has resulted in increasingly more papers citing sex-biased effects, the methodology is often flawed (11–13). Most studies do not disclose the biological sex of the cells or they pool samples from male and female donors without first adequately determining any sex-biased effects (11,14). Further, other studies lack enough statistical power to significantly determine the impact of biological sex beyond differences due to donor variability (13).

Human mesenchymal stromal cells (hMSCs) have the potential to promote repair and regeneration to address degenerative conditions or traumatic injury and are viable for cell therapy for a variety of tissues, including those in the musculoskeletal, neural, and immune systems (15–17). Additionally, hMSCs are a common target cell type for regeneration based on both their direct capacity for differentiation *in vitro* and their potent trophic effects *in vitro* and *in vivo* (18–20). However, despite significant evidence that hMSCs have the potential to modulate a large number of regenerative and inflammatory factors, consistent efficacy remains a challenge due to donor variability (21,22). Further, hMSC donor sex provides another layer of variability with significant differences in proliferative capacity, differentiation, and levels of senescence in human and non-human MSCs based on sex (22). One critical differentiator in donor sex is male and female baseline hormonal exposure and/or the difference in tissue and donor response to the same concentration of hormones (11). Despite this, traditional culture conditions for hMSCs contain exogenous E2 and E2-mimetic compounds that have an unknown effect on male and female hMSCs. The isolation of E2 exposure to male and female hMSCs allows us to better classify how sex differences contribute to variations in hMSC efficacy for cell therapy, which cannot be understood without a full and thorough accounting of E2 in cell culture media.

Phenol red and E2 in fetal bovine serum (FBS) are the two major sources of E2 or E2-mimetic compounds in traditional culture media (**Figure 1A**) (23–26). Phenol red is known to act in a similar manner to E2, however its full effects are not well understood. It has long been known that phenol red enhances the proliferative capacity of cells, particularly those that express high levels of estrogen receptors such as MCF-7 breast cancer cells (23), predominantly through ligand binding to the estrogen receptor alpha (ER-α) (23,25). Phenol red is present in cell culture media at levels much higher than traditional serum levels of E2 (23). While some attention has been given to this issue, previous results investigating phenol red were conducted solely in male cells or from non-human sources (27,28). Additionally, these previous studies only included a single donor, limiting the ability to statistically determine sex-biased effects apart from individual donor differences. Regarding E2 in FBS, many studies use charcoal dextran filtered FBS (CD-FBS) to remove E2 – among other constituents (29). However, what is known on the impact of CD-FBS on cell fundamental processes and phenotypes is limited. Previous work in osteoblasts found that the impacts of CD-FBS include altering metabolic profiles, proliferation, and differentiation (30). Previous work in bovine embryos found that CD-FBS supported increased embryonic viability through improved mitochondrial function by CD-FBS upregulation of SIRT1 (29). While FBS donor variability is known to effect hMSCs (31,32), no studies have directly investigated the effects of hormone-depleted FBS for hMSC culture in male and female cells. This issue also extends to the clinic, where the majority of clinical trials are utilizing some form of exogenous estrogens in their cell processing prior to treatment (**Supplementary Table 1**). Without systematic determination of culture media baseline levels of these E2-mimetic compounds, it is extremely difficult to decouple the impact of dosed E2 from the contributions of these exogenous estrogens.

**Figure 1.**
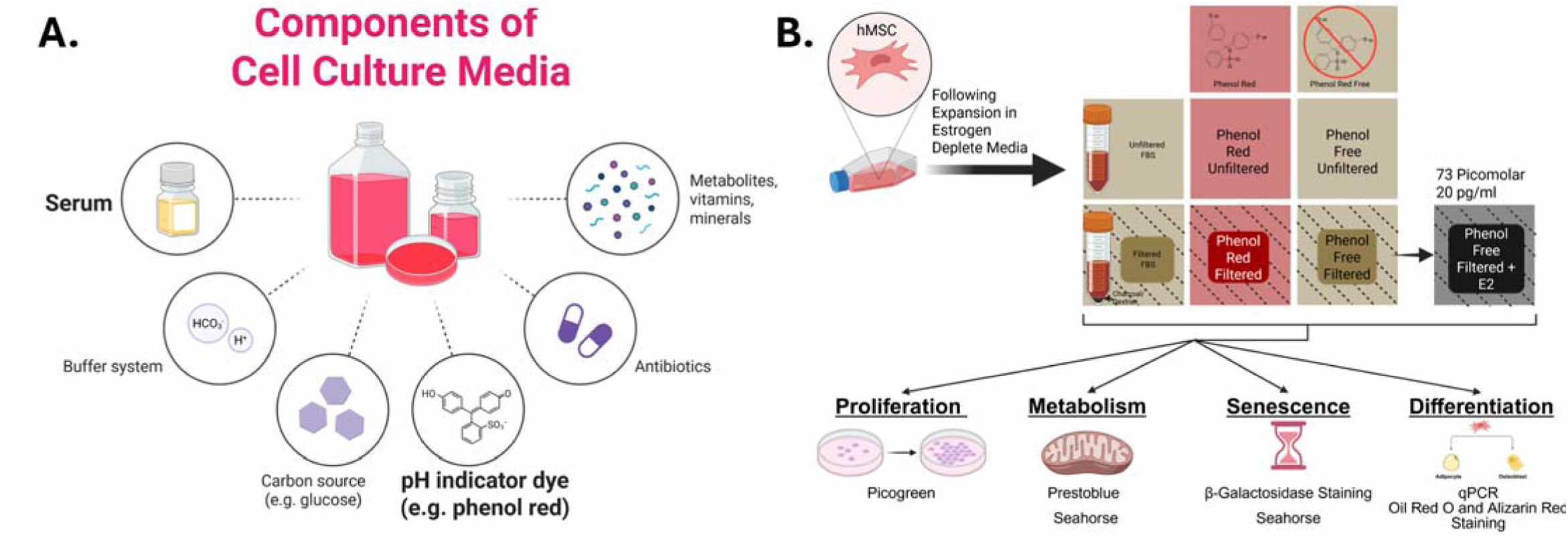
The common components of standard cell culture media highlighting serum and phenol red with estrogen-mimetic effects (A) Study design with color and shading scheme for media compositions (B). Schematics made in Biorender.

Herein, we investigate the effect of phenol red and unfiltered FBS on male and female hMSCs proliferation, metabolism, senescence, and adipogenic and osteogenic differentiation potential (**Figure 1B**). We utilized 8 donors, 4 male and 4 female, aged 19-23 and derived primarily from bone marrow. Proliferation and metabolic capacity were assessed via PicoGreen and PrestoBlue assays, respectively. Basal metabolic rate and senescence were measured utilizing a seahorse XF analyzer and β-galactosidase staining, respectively. Differentiation was assessed via key mRNA transcripts using quantitative polymerase chain reaction (qPCR) and differentiation capacity for osteogenic (alizarin red staining) and adipogenic (oil red o staining) lineages. This work provides a baseline for the effects of common exogenous estrogens in cell culture media and can be utilized by other researchers as a guide for determining which sources of exogenous estrogens are permissible for their given application. Overall, care must be taken when deriving data from male and female hMSCs in traditional culture conditions as their cellular health and phenotypic baseline may be biased by the media composition that is chosen.

## Methods

### Cell Culture

Cells were acquired from RoosterBio, ATCC, and StemBioSys with 4 male donors ranging from 19-22 years old and 4 female donors ranging from 20-23 years old (**Table 1**). hMSCs were harvested from bone marrow (n = 7) or adipose tissue (n = 1). Cells were cultured in phenol red-free, low glucose DMEM (Phenol Red: REF#11054-020, Phenol Free: REF#11885-084) supplemented with 10% FBS (Atlanta Biologics #S11650) and 4 mM L-glutamine (Gibco #21051024). Cells were expanded for 1 passage, plated at 9,000 cells/cm^2^, allowed to adhere for 24 hours, and then replaced with respective media conditions detailed below.

**Table 1:** hMSC Donor Details.

| Donor: Age (years), Sex | Source |
| --- | --- |
| 19, Male | RoosterBio, Bone Marrow |
| 21, Male | RoosterBio, Bone Marrow |
| 21, Male | StemBioSys, Bone Marrow |
| 22, Male | RoosterBio, Bone Marrow |
| 20, Female | RoosterBio, Bone Marrow |
| 21, Female | StemBioSys, Bone Marrow |
| 21, Female | Stemcell Tech, Adipose |
| 23, Female | ATCC, Bone Marrow |

### Media Formulations

Five different media formulations were made to test the effects of the estrogen-mimetic compound phenol red and E2 in FBS on hMSC properties. These consisted of the permutations of either DMEM with phenol red or phenol red free DMEM, and charcoal dextran filtered FBS (filtered FBS) or unfiltered FBS (normal FBS) (**Figure 1B)**. The most E2 depleted form of this media (phenol free, filtered FBS) was then supplemented with 71 picomolar 17β-estradiol (E2, Sigma E4389) to mimic the E2 levels assayed from the normal FBS per the supplier certificate of analysis (**Supplementary Image 1**).

### Picogreen

Cell proliferation was assessed by determining cell number using dsDNA quantification after five days in culture and relating back to the number of seeded cells. Cell membranes were disrupted and lysed with 1% Triton X-100 followed by pipetting up and down to access the internal dsDNA and allowed to lyse for 10 minutes. An Invitrogen™ Quant-iT™ PicoGreen™ dsDNA Assay Kit was utilized following manufacturer’s methods. Cell number was calculated assuming 7 picograms of DNA per cell (33).

### PrestoBlue

hMSC metabolism was assessed via a PrestoBlue assay to determine cell reductive capacity. Daily through 5 days, cell media was removed and replaced with 10% HS PrestoBlue reagent (Thermofisher PN# P50201) in phenol free, filtered FBS media for 1 hour. Following the incubation, fluorescence was read at 560 nm excitation/590 nm emission wavelengths in a Tecan Infinite 200 PRO plate reader followed by replenishment with fresh media of each respective treatment group. This assay is non-destructive and as such allows the same set of cells to be assayed across 5 days.

### □-galactosidase staining

Cells were fixed on day 5 and stained with Cell Signaling Technology Senescence β-Galactosidase Staining Kit (Cat#9860) following manufacturer’s methods for senescence associated signaling. Images of senescent associated staining were taken in a grid based on the center of the well with at least a 4×6 grid and summed to get the total number of positively staining residues per well, the same method was utilized on the same plate for DAPI nuclear staining. The number of senescent positive deposits are then divided by the number of nuclei to get the percentage of ‘activated’ cells. Analysis is done utilizing color deconvolution with custom color vectors to capture the maximum amount of blue staining, see **Supplementary Text 1** for code. Results are then checked by hand if they fall outside of ‘expected parameters’ I.E. an activation of >200% or extremely low or high counts for either deposits of senescent staining or DAPI. A mask was used from the deconvoluted red channel to remove imaging and staining artifacts such as crystallized stain.

### qPCR

Relative transcript levels of key osteogenic, adipogenic, and chondrogenic genes were assessed to provide evidence of media conditions impacting cell phenotype. For osteogenic differentiation, COL10a1 and RUNX2 were selected as COL10a1 is a core functional ECM protein for osteoblasts and RUNX2 is the main driver of chondrocyte hypertrophy to the osteochondral pathway (34). For adipogenic differentiation, PPARγ is a primary driver of adipogenic differentiation, and FABP4 is upregulated in adipogenesis and involved in fatty acid synthesis (35,36). Chondrogenic genes selected were SOX9 which is known to drive chondrogenic gene expression and ACAN, encoding for the protein core of the aggrecan proteoglycan and a major component of the cartilaginous extracellular matrix (37,38). Collagen type 1 was added to assess general extracellular matrix production and estrogen receptor alpha (ESR1) was run as it is the receptor target for phenol red (23). Cells were grown for 5 days in their respective media conditions without differentiation supplements then lysed according to manufacturer’s protocol (Thermofisher, PureLink™ RNA Mini Kit). TaqMan fast advanced master mix (Invitrogen #4444556) was used with RnaseOUT recombinant ribonuclease inhibitor (Invitrogen #10777019). Samples were eluted into 50 μL ultrapure water and reverse transcribed to cDNA using the Applied Biosystems high-capacity cDNA reverse transcription kit. Samples were analyzed utilizing Tapestation high-sensitivity (Agilent #5067-5579) RNA screen tape for representative RIN values and Qubit was run for RNA quantification. Samples with RIN values less than 7 were excluded. Qubit samples were run in duplicate with fresh RNA standards (Invitrogen #Q10210). Standard curves were performed on primers to determine the concentration where a linear response exists on a logarithmic scale for all reactions. cDNA libraries were diluted to 10 ng cDNA per reaction and assayed for GAPDH (Hs02786624_g1) (housekeeping gene), osteogenic targets RUNX2 (Hs01047973_m1) and Col10a1 (Hs00166657_m1), adipogenic targets PPARG (Hs01115513_m1) and FABP4 (Hs01086177_m1) chondrogenic targets Col1a1 (Hs00164004_m1), SOX9 (Hs00165814_m1), ACAN (Hs00153936_m1), and estrogen receptor alpha ESR1 (Hs01046816_m1). Samples were normalized by the delta-delta CT method to pooled reference cDNA from all hMSC donors cultured in phenol red unfiltered FBS.

### Differentiation Lineage Staining with Alizarin Red and Oil Red O

Trilineage differentiation was performed to determine if exogenous estrogens impacted functional differentiation throughout the process. For osteogenic lineage assessment, hMSCs from each respective media group were differentiated for 2 weeks in osteogenic media containing 100 nM Dexamethasone (Thermo Scientific Chemicals# A17590.03), 10 mM Beta-Glycerophosphate (Thermo Scientific Chemicals# L03425.36), and 50 ug/ml Ascorbic Acid (Sigma-Aldritch# 49752). For adipogenic assessment, hMSCs were differentiated for 2 weeks in adipogenic media containing 1 uM Dexamethasone (Thermo Scientific Chemicals # A17590.03), 500 uM Isobutylmethylxanthine (Thermo Scientific Chemicals # J64598.MC), 5 ug/ml Insulin (Gibco# 12585014), and 200 ug/ml of Ascorbic Acid (Sigma-Aldritch# 49752). Image analysis was performed utilizing specifically designed ImageJ macros (39). For alizarin red staining, the images were blurred with a Gaussian filter (sigma = 2) to reduce noise. Then a custom color deconvolution process was employed to pull out the specific color of positively stained calcium deposits. “Analyze particles” was then used to determine the number of positively stained calcium deposits with a size gate (1000 pixels – infinity) to prevent stain particulates crashing out of solution from being counted. See **Supplementary Text 2** for example code. For oil red o analysis, the images were blurred with a Gaussian filter (sigma = 5) to distinguish between signal and remove extremely small particles followed by a custom color deconvolution process to pull out the specific color of positively stained lipid droplets. This color profile was determined by averaging the color vectors of clearly positively stained lipid droplets. Each image is then run through auto-local threshold before being watershed to isolate individual droplets and counted using “analyze particles” with a circularity gate to prevent off target staining from being counted. See **Supplementary Text 3** for example code.

### Statistical Analysis

Skewness of data was verified in R as being less than 0.5, which according to Webster and Oliver is not enough to significantly alter the normality of the dataset (40–42). Statistical comparisons on the graph are done through Bonferroni multiple comparisons testing and 2-way ANOVA. Data was also fit to a linear model in R Studio (43–48) for each measured outcome to better capture overall variability and explain the contribution of each source of exogenous estrogen as well as donor sex. Akaike information criterion (AIC) values were used to determine the optimal interaction parameters for each model (49). A summary of the P values for each parameter is listed in **Table 2**. Each model was run to test for interaction between phenol red, unfiltered FBS, and E2 supplementation individually and combined.

**Table 2:**
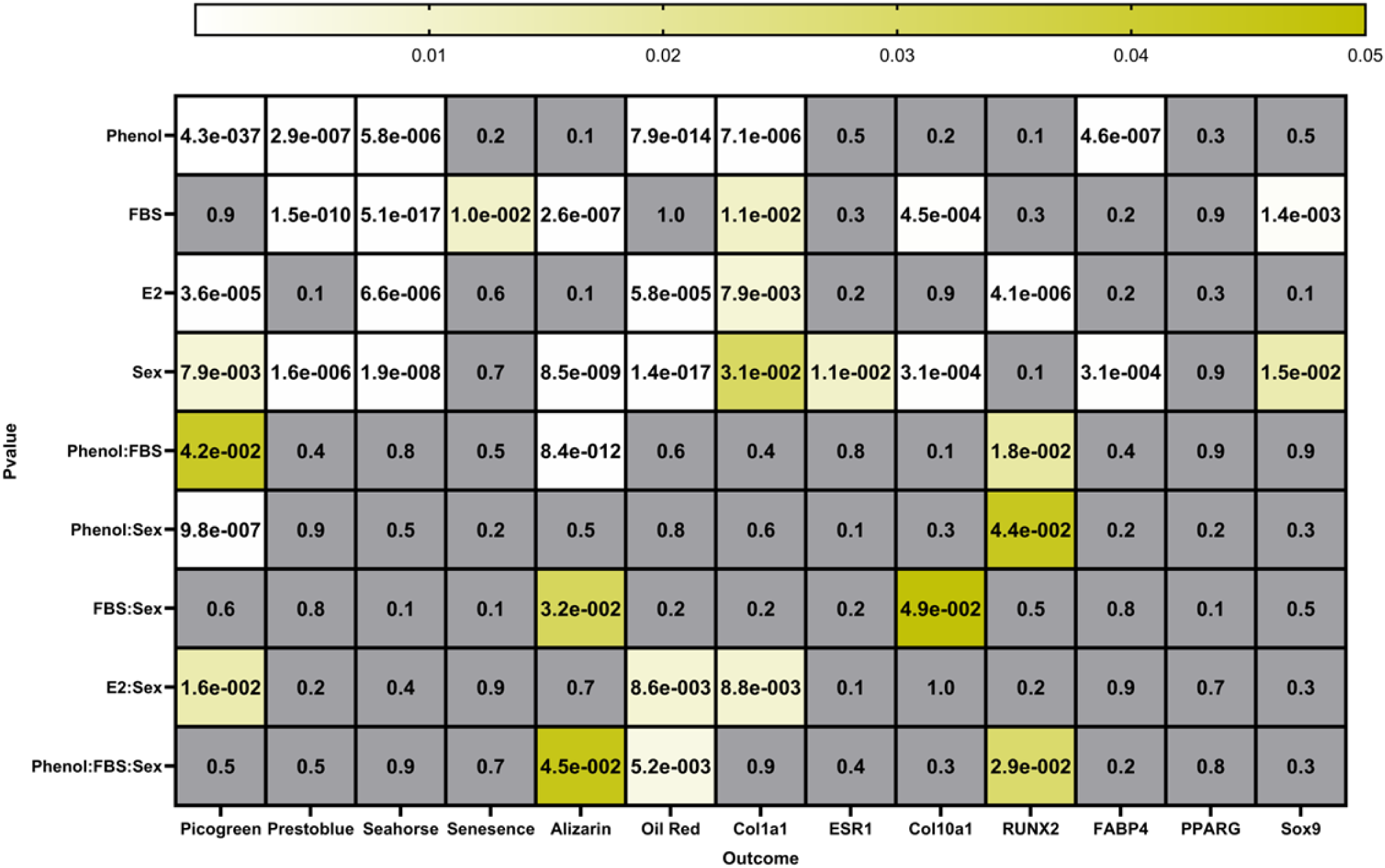
Linear Model Parameters to show impact and interaction of culture conditions and donor sex. Values greater than 0.05 are in grey as they are insignificant. Values less than 0.05 are shown on a gradient from less significant (yellow, 0.05) to more significant (white, 4.3*10^-37).

## Results

For all outcomes, compiled results from all donors are shown in the main figures and individual donor graphs are displayed in **Supplementary Figures 2-14**.

### Proliferation

Phenol red-containing media was the primary driver of cell proliferation across all donors **(Figure 2A**). Overall, female cells proliferated significantly more in the presence of phenol red compared to male cells (**Figure 2B**), an effect not observed in phenol free conditions (**Figure 2B**). E2 supplementation of the phenol free, filtered FBS group suppressed growth in female cells, but not in male cells when compared to the phenol free, filtered FBS group on its own (**Figure 2A)**. However, this effect was not significantly different when directly comparing the impact of E2 between male and female hMSCs (**Figure 2B**). Filtered vs. normal FBS did not significantly impact proliferation across all donors (**Figure 2A)**. From linear model analysis, the interaction between sex and phenol red treatment was highly significant, indicating that sex significantly impacts the response of these cells to phenol red in a manner beyond which is explained simply by the sex differences alone without phenol red in the media. Significant interactions were also observed between phenol red and FBS, indicating FBS alters the capacity of phenol red to affect proliferation.

**Figure 2.**
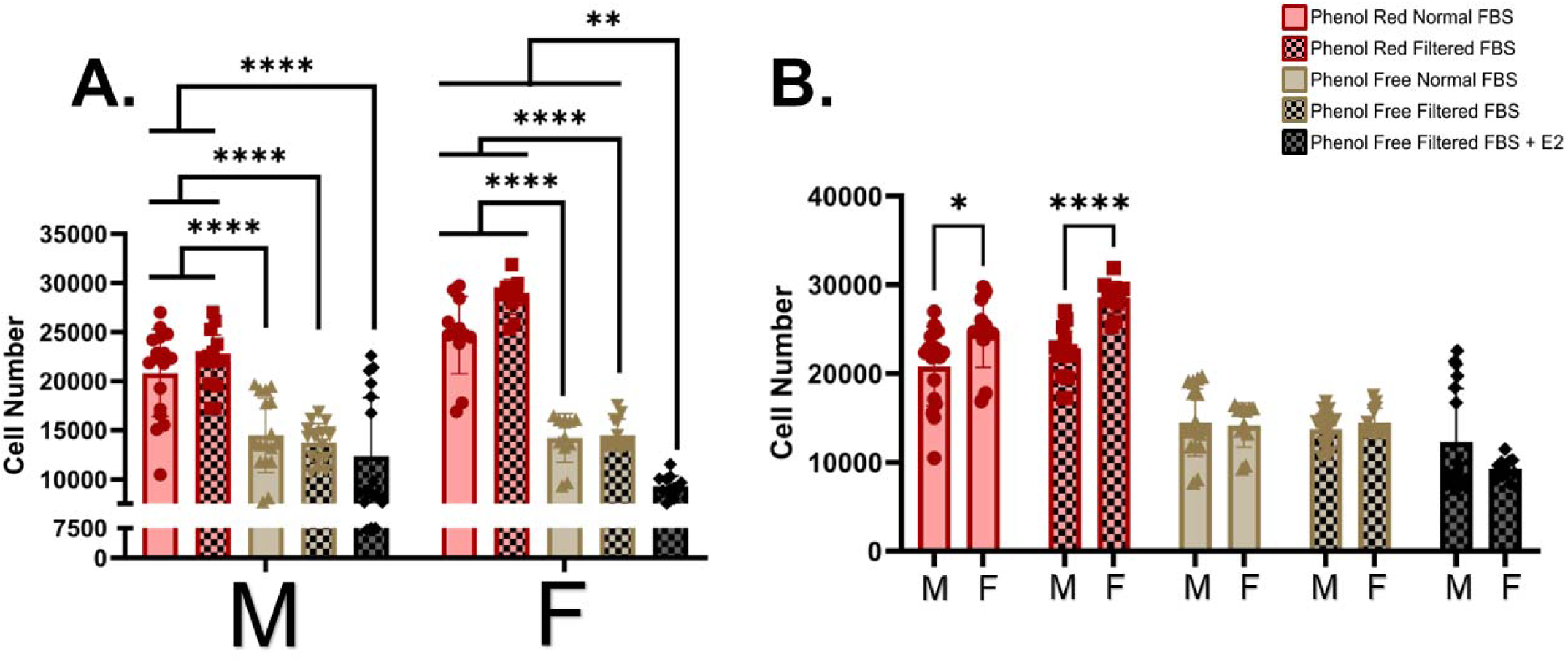
hMSC cell numbers at Day 5 from PicoGreen dsDNA assessment to determine media impacts on proliferation. Statistical comparisons done within donor sex (A) Statistical comparisons done between donor sexes (B). All data tested for normality utilizing the Shapiro-Wilk test (α = .05) and analyzed utilizing 2-way ANOVA and Bonferroni multiple comparison testing. Picogreen Linear Model statistics Phenol: P=4.32e^-37^, FBS: P=0.934, E2: P=3.64e^-5^, Sex: P=0.008 Interactions: Phenol:FBS: P=0.042, Phenol:Sex: P=9.78e^-7^, FBS:Sex: P=0.565, E2:Sex: P=0.016, Phenol:FBS:Sex: P=0.504 P<0.0001. * P =< .05, ** P =< .01, *** P =< .001, **** P =< .0001. Each dot represents a different well in a tissue culture plate: 3 wells per donor X 4 donors = 12 points per condition. N represents biological replicate of individual donors (3 replicates per donor)

### Metabolism

The PrestoBlue assay was used to measure the hMSC populations’ capacity to convert resazurin to resorufin, an indicator of cells’ ability to covert NADPH to NADP (50), through 5 days of culture in each respective media. FBS filtration was the major driver of changes in metabolic activity with reduced metabolic activity of hMSCs in both male and female cells in the presence of phenol red (**Figure 3A**). In female cells, the metabolic differences with filtered FBS were non-significant in phenol red free media; however, this could be driven by an individual female donor that was particularly metabolically active (**Figure 3A and Supplemental Figure 3)**. Low level E2 supplementation (71 pM) did not recover metabolic activity in hMSCs (**Figure 3A**). Female hMSCs had significantly higher metabolic activity than males in the phenol free filtered FBS media (**Figure 3B**). PrestoBlue time course plots demonstrate that female cells exhibited a higher capacity for metabolic activity compared to males (**Figure 3C and 3D**). Linear modelling indicated significant effects from sex, phenol red, FBS, and E2 on the reductive capacity of male and female cells. However, the interaction parameters were non-significant, indicating that the effect of exogenous E2 on the reductive capacity of hMSCs is mostly explained by either baseline sex differences or the levels of exogenous E2 in the media not by a sex specific response to E2.

**Figure 3.**
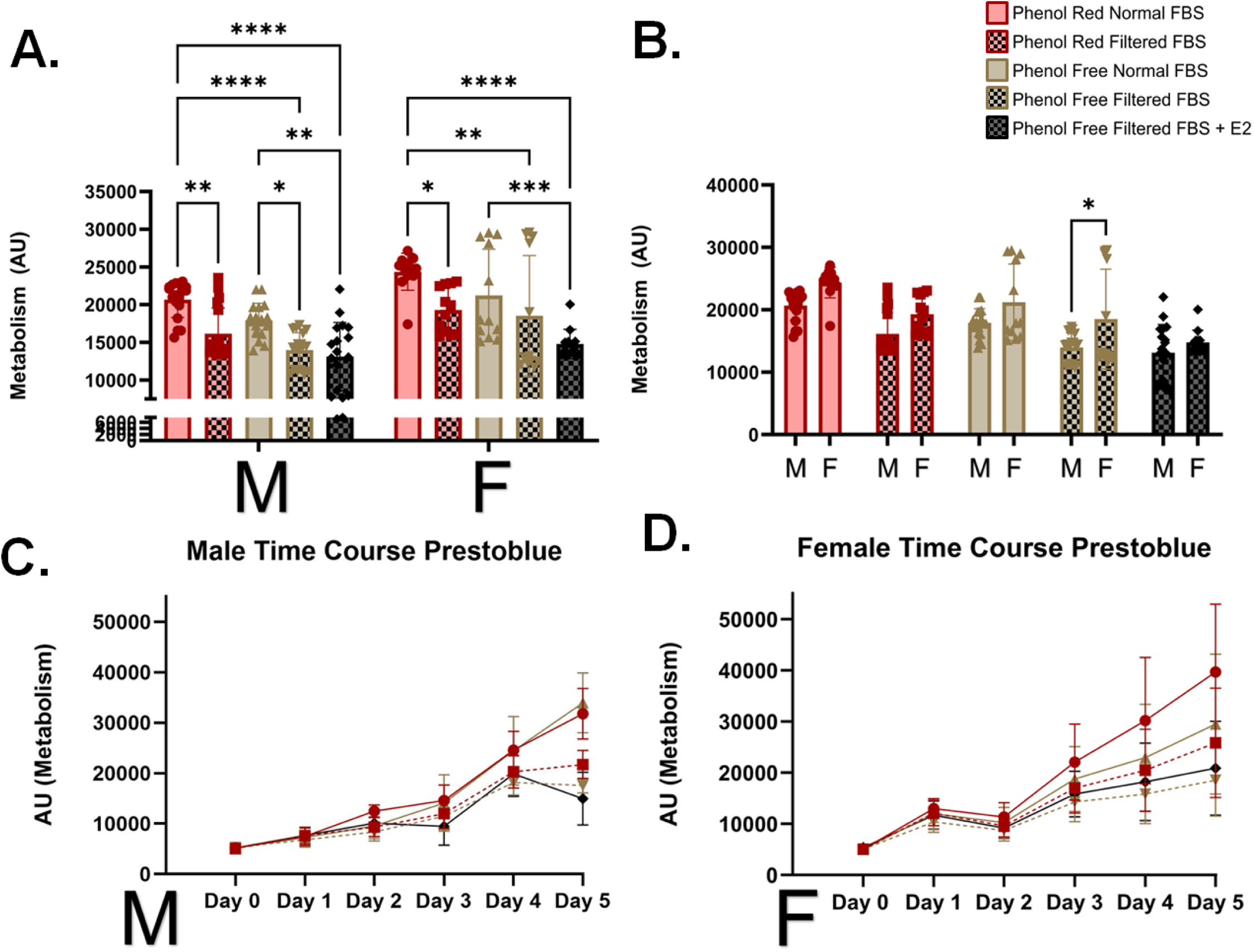
hMSC PrestoBlue metabolism through Day 5. Statistical comparisons done within donor sex (A). Statistical comparisons done between donor sexes (B). Male time course metabolism data through Day 5 (C). Female time course metabolism data through Day 5 (D). All data tested for normality utilizing the Shapiro-Wilk test (α = .05) and analyzed utilizing linear model analysis. Prestoblue Linear Model statistics Phenol: P=2.85e^-7^, FBS: P=1.49e^-10^, E2: P=0.076, Sex: P=1.58e^-6^ Interactions: Phenol:FBS: P=0.370, Phenol:Sex: P=0.870, FBS:Sex: P=0.800, E2:Sex: P=0.211, Phenol:FBS:Sex: P=0.545* P =< .05, ** P =< .01, *** P =< .001, **** P =< .0001. Each dot represents a different well in a tissue culture plate: 3 wells per donor X 4 donors = 12 points per condition.

Because of this stark shift in metabolic profile, a deeper metabolic profile utilizing the glycolysis stress test kit on the Seahorse XF Analyzer was done. Interestingly, the basal metabolic profile, namely the ratio of the oxygen consumption rate (OCR, indicative of oxidative phosphorylation) to extracellular acidification rate (ECAR, indicative of glycolysis), was significantly higher only in female cells in response to unfiltered FBS **(Figure 4A**). Specifically, normal FBS caused an increase in the prevalence of oxidative phosphorylation compared to glycolysis in female hMSCs (**Figure 4A**). Female donors had a significantly lower OCR/ECAR in phenol red filtered FBS when compared to matched male donors and significantly higher OCR/ECAR in phenol free normal FBS indicating their sensitivity to exogenous E2 in FBS (**Figure 4B**). Low-level E2 supplementation did not promote a significantly higher OCR/ECAR metabolic balance, indicating this mechanism may not be solely mediated by pM levels of E2. Based on comparisons between donor sex, sex does not interact with any treatment to significantly affect metabolism, rather there are baseline differences in metabolism based on sex that are altered by phenol red, FBS, and E2 indicating that male and female donors have sex specific responses to the exogenous forms of E2 when it comes to their basal metabolic rate.

**Figure 4.**
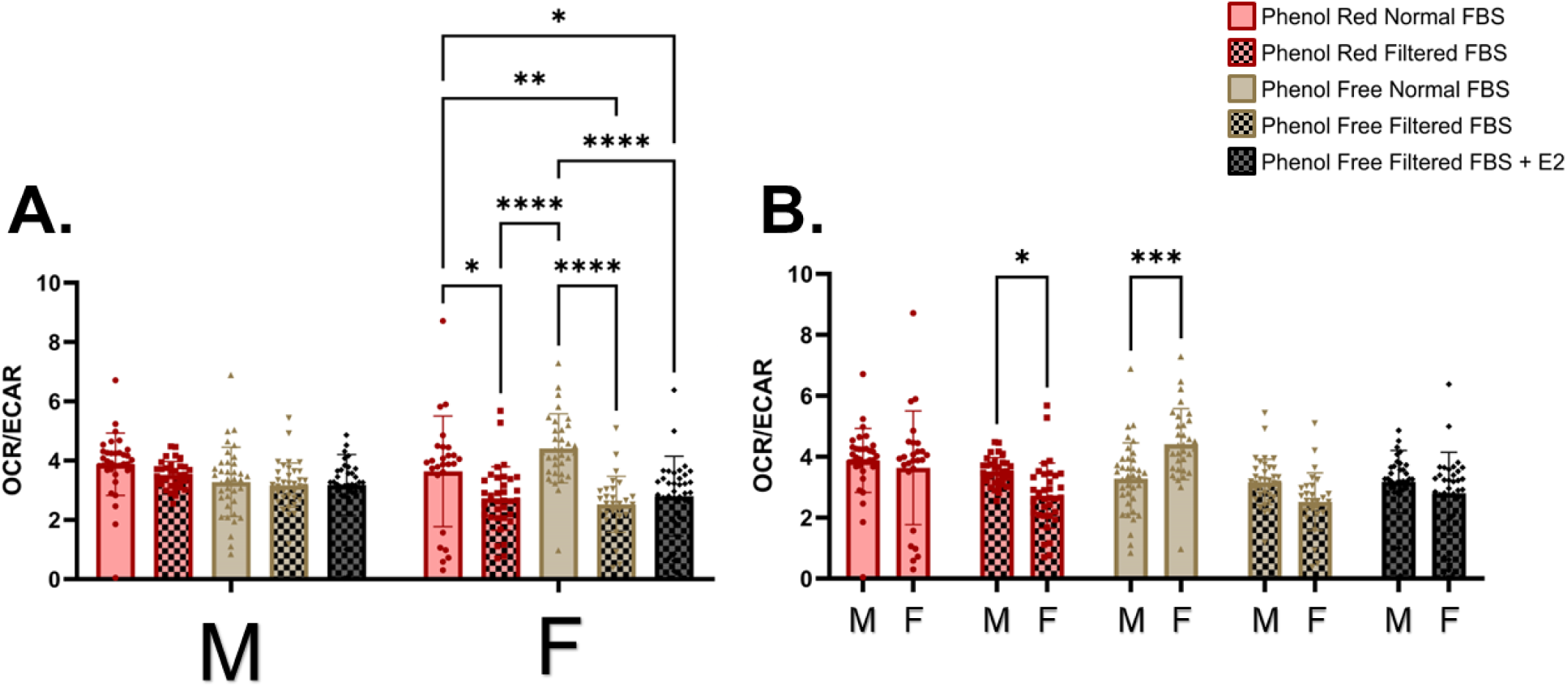
Seahorse Basal Metabolic Rate Data on Day 5. Statistical comparisons done within donor sex (A) Statistical comparisons done between donor sexes(B). All data tested for normality utilizing the Shapiro-Wilk test (α = .05) and analyzed utilizing linear model analysis. Seahorse Linear Model statistics Phenol: P=5.78e^-6^, FBS: P=5.10e^-17^, E2: P=6.60e^-6^, Sex: P=1.89e^-8^ Interactions: Phenol:FBS: P=0.849, Phenol:Sex: P=0.480, FBS:Sex: P=0.101, E2:Sex: P=0.407, Phenol:FBS:Sex: P=0.896* P =< .05, ** P =< .01, *** P =< .001, **** P =< .0001. Each dot represents a different well in a tissue culture plate, 3 wells per donor, 4 donors, for a total of 12 points per condition.

### Senescence

β-galactosidase staining was conducted to determine whether the media impact on proliferation was due to an increase in senescence. Overall, senescence-associated β-galactosidase staining levels were variable between donors (**Figure 5 and Supplementary Figure 4**). Senescence-associated beta-galactosidase staining indicated a similar pattern to that of the basal metabolic rate. Normal FBS significantly increased senescence-associated β-galactosidase staining in male donors (**Figure 5A)**. In contrast to the basal metabolic rate data, normal FBS also increased the senescence-associated β-galactosidase staining in the phenol red, but not the phenol free groups in the male cells. Low level E2 supplementation did not significantly alter senescence-associated staining. A difference between male and female cells was not observed, however individual donors behaved in distinct manners (**Supplementary Figure 5)** indicating a characteristic other than sex is driving changes. Normal unfiltered FBS was the main driver of senescence-associated staining and the only significant contributor to the variance when analyzed via a linear model.

**Figure 5.**
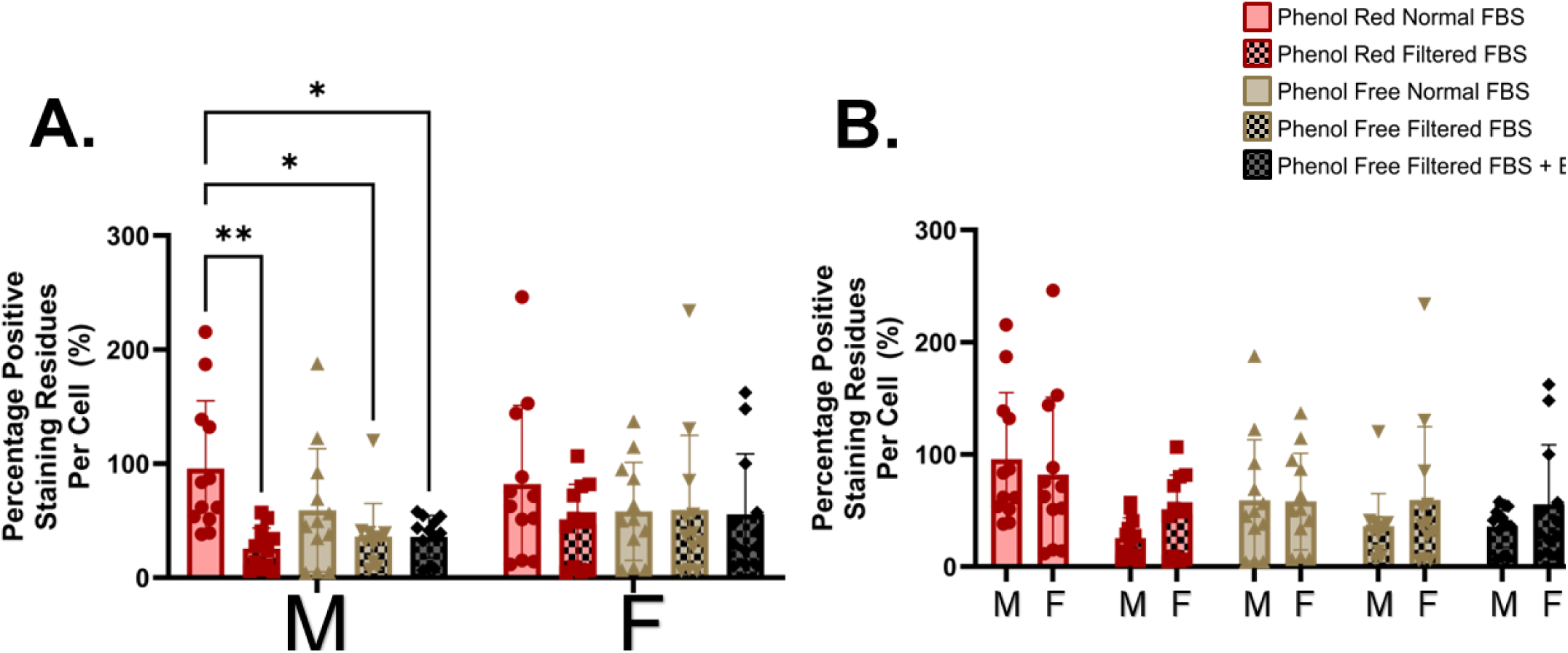
β-galactosidase-associated senescence staining on Day 5. Statistical comparisons done within donor sex (A) Statistical comparisons done between donor sexes. (B) Analyzed utilizing 2-way ANOVA and Bonferroni multiple comparison testing. ANOVA Statistics Interaction: P=0.5853, Row Factor (Sex): P<0.2577, Column Factor (Treatment): P<0.0027. * P =< .05, ** P =< .01, *** P =< .001, **** P =< .0001. Each dot represents a different well in a tissue culture plate, 3 wells per donor, 4 donors, for a total of 12 points per condition.

### Differentiation

To assess the impacts of exogenous estrogens in the media on the differentiation potential of male and female hMSCs, cells were differentiated in osteogenic and adipogenic media with their respective media differences (I.E. with or without phenol red and with either normal or unfiltered FBS) for 2 weeks and stained with alizarin red and oil red o to assess calcium deposits and lipid droplets, respectively. Images illustrating minimal, representative, and maximum staining are shown in **Figure 6A** and **6B**. Staining for alizarin red (**Figure 6)** revealed a decrease in osteogenic differentiation with unfiltered FBS when compared to filtered FBS in the presence of phenol red in both male and female hMSCs. Additionally, in female donors, the E2 supplemented condition was significantly lower in calcium deposition staining than the phenol free filtered FBS condition, indicating that in female cells picomolar E2 supplementation inhibits osteogenic differentiation. Female cells overall had more calcium staining in phenol red normal FBS, and both phenol free conditions, which matched to an overall presence of baseline sex differences. Response to FBS and sex was significant, while the effects of phenol red alone and E2 alone were not. Interaction between FBS and phenol red was highly significant. Additionally, the impact of the combination of phenol, FBS, and sex was significant, indicating all these factors contribute to differences in osteogenic differentiation. Overall, osteogenic differentiation was increased by normal FBS in phenol red containing media for both male and female donors, indicating higher levels of E2 and E2 mimetic compounds in media may accelerate osteogenic differentiation through day 14.

**Figure 6.**
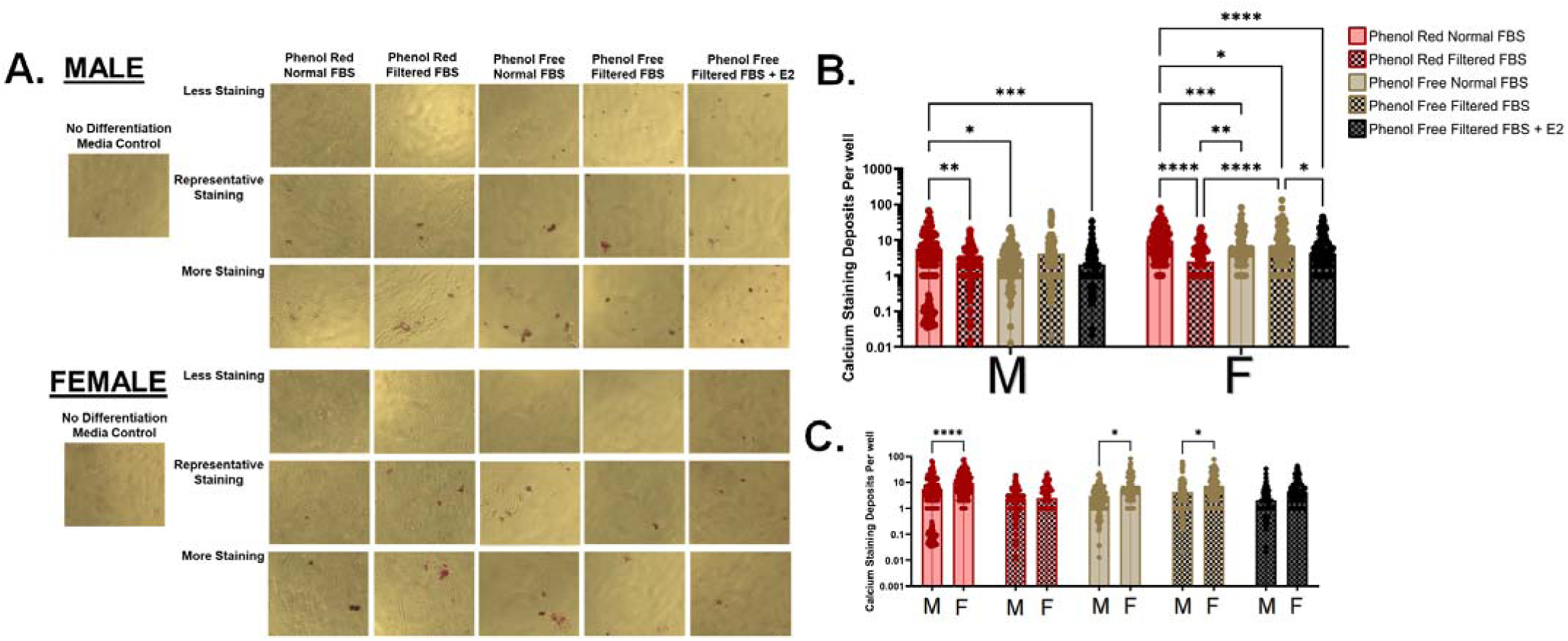
Alizarin Red staining to assess media impacts on osteogenic differentiation. Representative images ranging from less to more staining for each condition in male and female hMSCs. (A) Statistical comparisons done within donor sex. (B) Statistical comparisons done between donor sexes. (C) Analyzed utilizing linear model analysis. Alizarin Red Linear Model statistics Phenol: P=0.122, FBS: P=2.60e^-7^, E2: P=0.149, Sex: P=8.48e^-9^ Interactions: Phenol:FBS: P=8.43e^-12^, Phenol:Sex: P=0.522, FBS:Sex: P=0.032, E2:Sex: P=0.682, Phenol:FBS:Sex: P=0.045. * P =< .05, ** P =< .01, *** P =< .001, **** P =< .0001. Each point represents an image analyzed from a well, 3 wells per condition per donor.

Images revealing minimal, representative, and maximal oil red o staining for each media group from male and female donors are shown in **Figure 7A**. In male cells, phenol red conditions promoted significantly more adipogenic differentiation compared to phenol free conditions (**Figure 7A and 7B**). In female cells, loss of phenol red from the media reduced adipogenic differentiation in the unfiltered FBS condition, and this was not recovered by pM E2 supplementation of the phenol free unfiltered FBS condition. Female cells had higher amounts of adipogenic staining in phenol red conditions as well as the phenol free filtered conditions than male cells. The fact that the phenol free unfiltered FBS condition and E2 supplementation condition trended similarly indicates that this effect may be largely due to the pM presence of E2, and the higher amounts in phenol red containing media indicates that phenol red may be rescuing the adipogenic potential of these cells. Linear modelling indicates that phenol red, sex, and E2 supplementation are significant in altering adipogenic staining. Sex synergistically alters adipogenic staining but only with all other parameters interacting.

**Figure 7.**
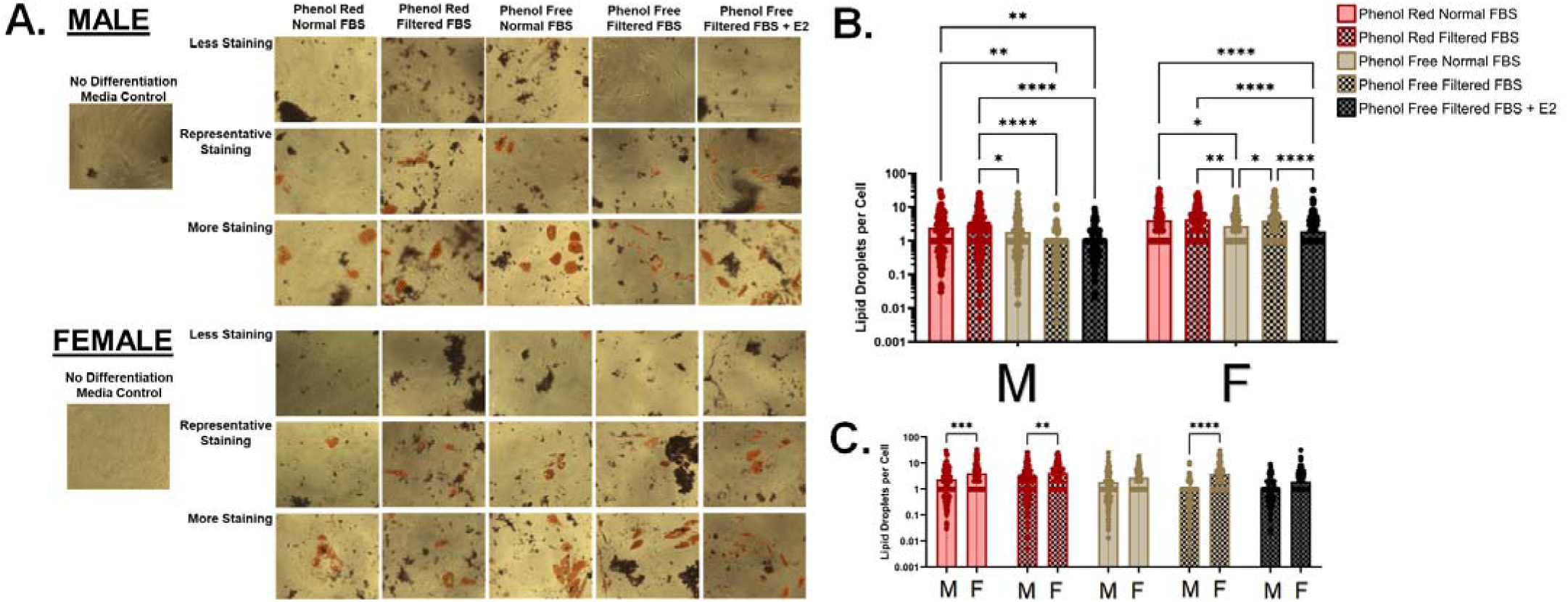
Oil Red O staining to assess media impacts on adipogenic differentiation. Representative images ranging from less to more staining for each condition in male and female hMSCs. (A) Statistical comparisons done within donor sex. (B) Statistical comparisons done between donor sexes. (C) Analyzed utilizing linear model analysis. Oil Red O Linear Model statistics Phenol: P=7.78e^-14^, FBS: P=0.968, E2: P=5.82e^-5^, Sex: P=1.40e^-17^ Interactions: Phenol:FBS: P=0.588, Phenol:Sex: P=0.820, FBS:Sex: P=0.243, E2:Sex: P=0.009, Phenol:FBS:Sex: P=0.005* P =< .05, ** P =< .01, *** P =< .001, **** P =< .0001. Each point represents an image analyzed from a well, 3 wells per condition per donor.

qPCR analysis was performed for 2 key genes −1 transcription factor, 1 downstream protein - for each tri-lineage path based on literature recommendations (51). Additionally, type 1 collagen (COL1a1) and estrogen receptor alpha (ESR1) were targets to investigate overall matrix deposition alterations and the activity of the estrogen receptor that phenol red is known to bind. Cells for qPCR assessment were cultured without differentiation media such that significant differences in mRNA expression of any target gene are solely due to the differences in the media conditions of interest in this study rather than compounds in the differentiation media. Osteogenic differentiation was assessed by RUNX2 and Col10a1 expression, adipogenic differentiation was assessed by FABP4 and PPARγ, and chondrogenic differentiation was assessed by SOX9 and Col2a1. Col2a1 failed to amplify in any of the tested conditions and was therefore excluded from the final data.

Type 1 collagen expression was not significantly altered between individual groups except E2 supplementation in male cells (**Figure 8 A and B)**. Linear modelling revealed that type 1 collagen was significantly altered by phenol red, CD-FBS, donor sex, and E2 supplementation despite the lack of significantly different direct comparisons. Additionally, the expression of COL1a1 in response to E2 supplementation was significantly higher in male cells, indicated by the significant interaction between E2 supplementation and donor sex as well as the direct comparison. Cells appear to express more COL1a1 in exogenous E2 deplete conditions, but not significantly so.

**Figure 8.**
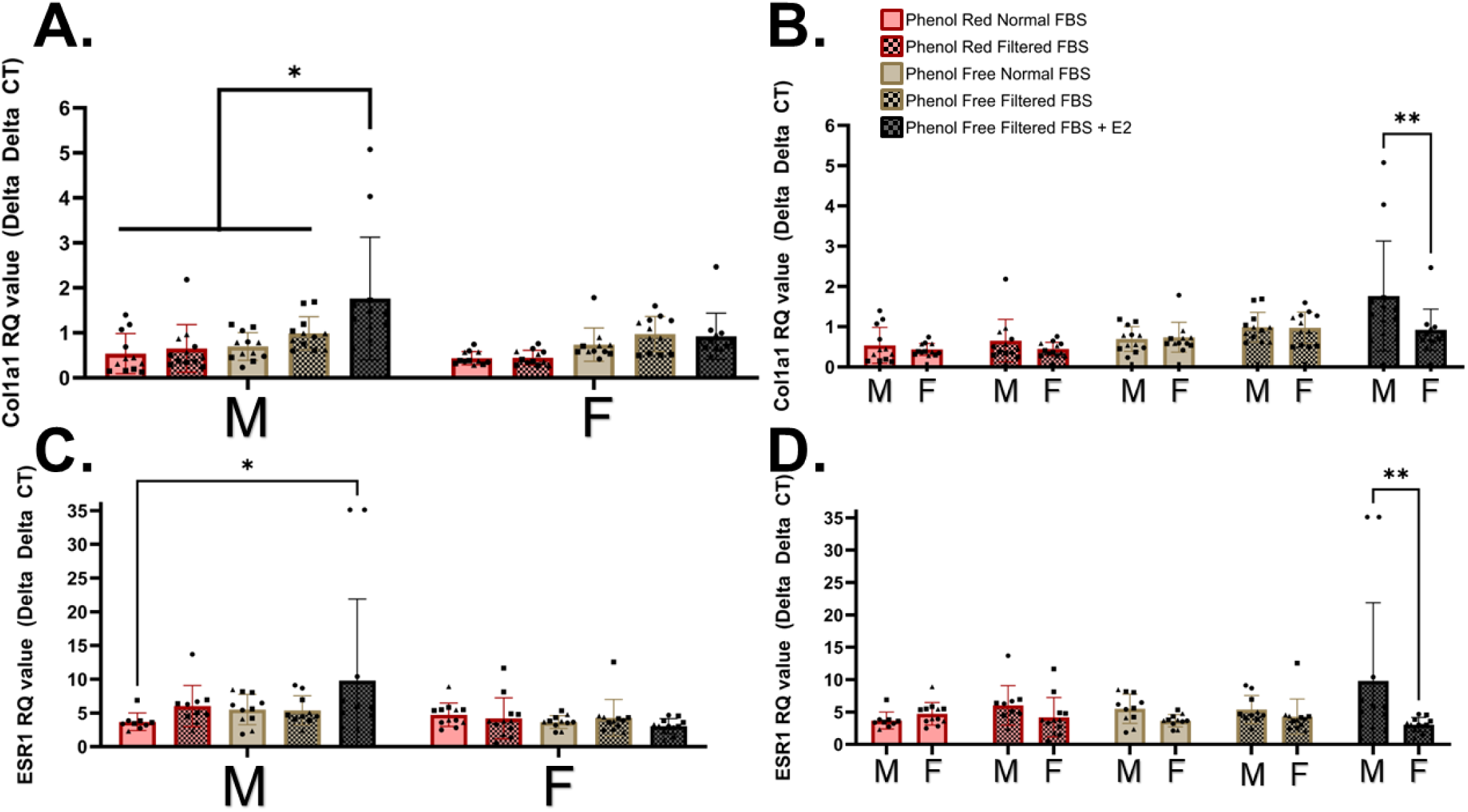
Col1a1 and ESR1 expression. Statistical comparisons done within donor sex for Col1a1 (A) and ESR1 (C). Statistical comparisons done between donor sexes for Col1a1 (B) and ESR1 (D). Analyzed utilizing linear model testing. **Col1a1** Linear Model Statistics Phenol: P=7.10e^-6^, FBS: P=0.011, E2: P=0.008. Sex: P=0.031 Interactions: Phenol:FBS: P=0.375, Phenol:Sex: P=0.564, FBS:Sex: P=0.152, E2:Sex: P=0.009, Phenol:FBS;Sex: P=0.922 **ESR1** Linear Model Statistics Phenol: P=0.474, FBS: P=0.304, E2: P=0.224. Sex: 0.011. Interactions: Phenol:FBS: P=0.834, Phenol:Sex: P=0.101, FBS:Sex: P=0.167, E2:Sex: P=0.059, Phenol:FBS;Sex: P=0.352 * P =< .05, ** P =< .01, *** P =< .001, **** P =< .0001. Each dot represents a different well in a tissue culture plate, 4 wells per donor, 3 donors per sex, for a total of 12 points per condition.

Estrogen receptor alpha expression was upregulated in male cells in response to E2 supplementation of the phenol free, filtered FBS media compared to female cells, although this is largely from the contribution of a single male donor (**Figure 8 C and D, Supplementary Figure 8).** The upregulation in male estrogen receptor alpha expression in response to phenol free, filtered FBS supplemented with E2 was also significantly higher than all other media conditions for male cells. Interestingly, the only significant differences in E2 expression were inherent differences based on sex, according to the linear model analysis.

Female cells had a significantly lower expression of RUNX2 in phenol red filtered FBS conditions. RUNX2 in the phenol free filtered FBS condition was significantly upregulated in female cells when compared to other filtered FBS conditions, with a similar trend in male cells. E2 supplementation suppressed RUNX2 expression in both male and female cells (**Figure 9A and 9B**). In female cells, this effect was expanded with the E2 deplete condition showing significantly higher RUNX2 expression than both other filtered FBS conditions. Linear modelling of RUNX2 expression revealed that treatment with exogenous E2 and the interaction of donor sex with that treatment was significant, but the baseline differences between male and female donors was not. RUNX2 expression was significantly impacted primarily by E2 supplementation and interaction between phenol red, FBS condition, and donor sex. Phenol and FBS significantly altered RUNX2 expression together, but not alone. Additionally, phenol red, FBS, and donor sex all significantly interacted to alter RUNX2 expression indicating that osteogenic signaling may be impacted by all these factors.

**Figure 9.**
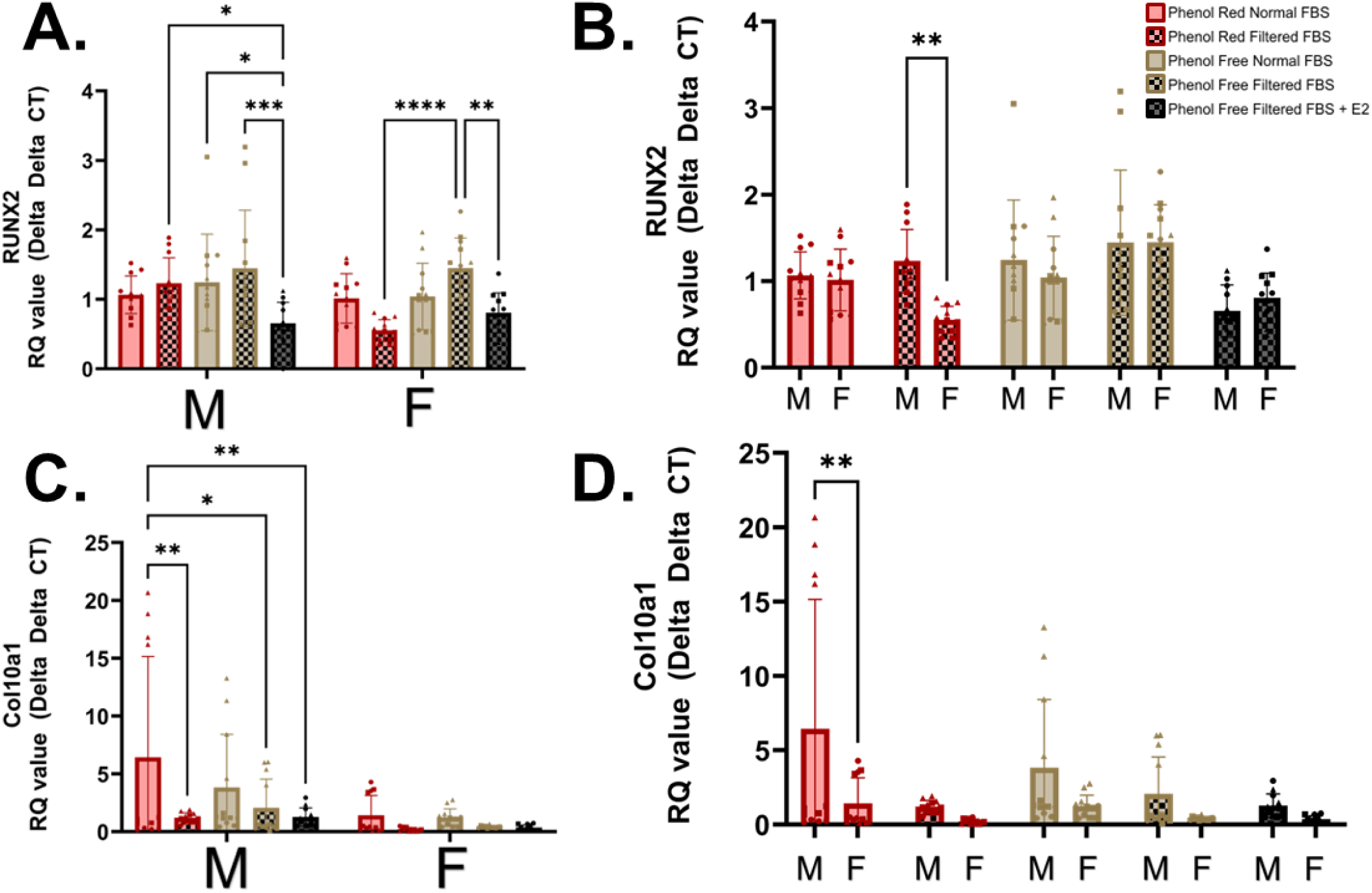
Osteogenic markers Runx2 and Col10a1. Statistical comparisons done within donor sex for Runx2 (A) and Col10a1 (C). Statistical comparisons done between donor sexes for Runx2 (B) and Col10a1 (D). RUNX2 Linear Model statistics Phenol: P=0.106, FBS: P=0.283, E2: P=4.14e^-6^, Sex: P=0.067 Interactions: Phenol:FBS: P=0.018, Phenol:Sex: P=0.044, FBS:Sex: P=0.537, E2:Sex: P=0.157, Phenol:FBS:Sex: P=0.029; Col10a1 Linear Model statistics Phenol: P=0.199, FBS: P=4.53e^-4^, E2: P=0.947, Sex: P=3.13e^-4^ Interactions: Phenol:FBS: P=0.148, Phenol:Sex: P=0.293, FBS:Sex: P=0.049, E2:Sex: P=0.983, Phenol:FBS:Sex: P=0.256. * P =< .05, ** P =< .01, *** P =< .001, **** P =< .0001. Each dot represents a different well in a tissue culture plate, 4 wells per donor, 3 donors, for a total of 12 points per condition.

Interestingly, the Col10a1 expression pattern did not match the expression pattern of its matched transcription factor RUNX2 (**Figure 9C and 9D**). Col10a1 expression was significantly higher in the male cells in the phenol red normal FBS condition than all the filtered FBS conditions. Expression in female donors was low and non-significant between groups – albeit the trend was similar to the male cells. Male cells in phenol red normal FBS media had significantly higher Col10a1 expression than female cells and females trended low for all other media conditions compared to males. Sex and exogenous E2 in the media did significantly contribute to the variance via linear model statistics. Most interactions were non-significant, however FBS did significantly interact with donor sex.

FABP4 expression (**Figure 10A and 10B**) showed a significant increase in the phenol red filtered FBS conditions, in agreement with the semi-quantification of oil red o staining, when compared to the phenol free unfiltered + E2 condition. Male cells additionally had significantly higher expression of FABP4 in both phenol red conditions, indicating phenol red may pre-prime hMSCs for adipogenic differentiation. Female donors in the phenol red normal FBS conditions had significantly lower expression of FABP4 compared to male cells. Linear modelling indicated significant differences based on sex and phenol red, but not their interaction, indicating that male cells may simply have a higher baseline for FAPB4 expression *in vitro*.

**Figure 10.**
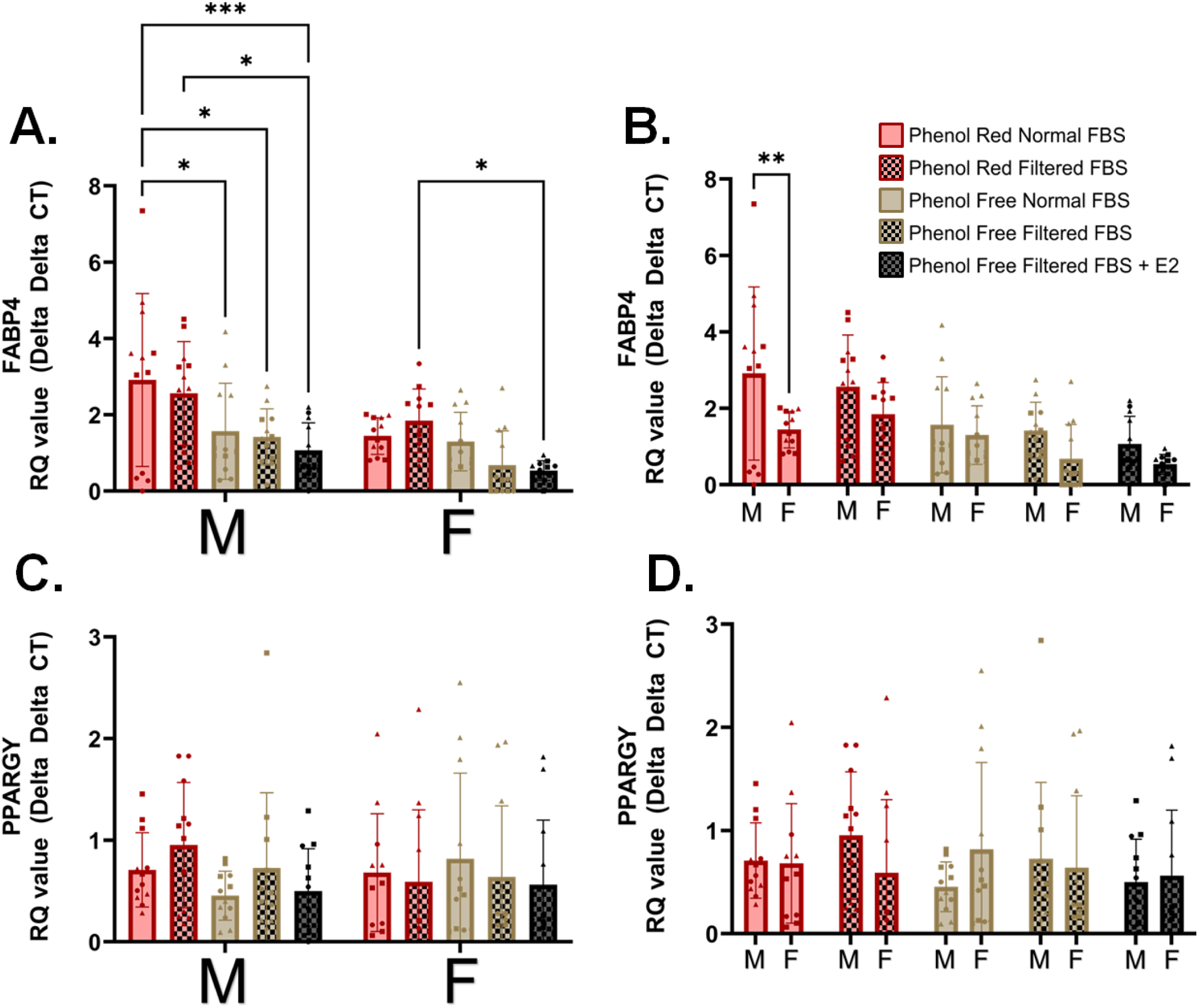
Adipogenic markers FABP4 and PPARG. Statistical comparisons done within donor sex for FABP4 (A) and PPARGY (C). Statistical comparisons done between donor sexes for FABP4 (B) and PPARGY (D). Analyzed utilizing linear model analysis. FABP4 Linear Model statistics Phenol: P=4.61e^-7^, FBS: P=0.180, E2: P=240, Sex: P=3.09e^-4^ Interactions: Phenol:FBS: P=0.363, Phenol:Sex: P=0.158, FBS:Sex: P=0.788, E2:Sex: P=0.870, Phenol:FBS:Sex: P=0.173 PPARG Linear Model statistics Phenol: P=0.309, FBS: P=0.893, E2: P=0.336, Sex: P=0.924 Interactions: Phenol:FBS: P=0.905, Phenol:Sex: P=0.179, FBS:Sex: P=0.123, E2:Sex: P=0.712, Phenol:FBS:Sex: P=0.815; PPARG ANOVA Statistics Interaction: P=0.3563, Row Factor (Sex): P=0.9237, Column Factor (Treatment): P=0.7332. * P =< .05, ** P =< .01, *** P =< .001, **** P =< .0001. Each dot represents a different well in a tissue culture plate, 4 wells per donor, 3 donors, for a total of 12 points per condition.

PPARG showed no significant differences in treatment or sex (**Figure 10C and 10D**). Individual donors did exhibit significant changes between the treatment conditions (**Supplemental Figure 13**); however, these effects are not seen when data is aggregated by sex, indicating that PPARG response to exogenous E2 may be donor dependent, and not sex dependent.

qPCR for SOX9 did not reveal any statistically significant direct comparisons **(Figure 11A and B)**. SOX9 expression did trend up with E2 supplementation, but the effect was not significant. Linear modelling indicated that both sex and the filtered FBS significantly impacted the variance. There were no significant interaction parameters, indicating sex differences would be largely due to baseline expression differences. Individual donor expression patterns were significant, with two female donors indicating SOX9 was maximized by phenol free, filtered FBS conditions (**Supplemental Figure 14)**.

**Figure 11.**
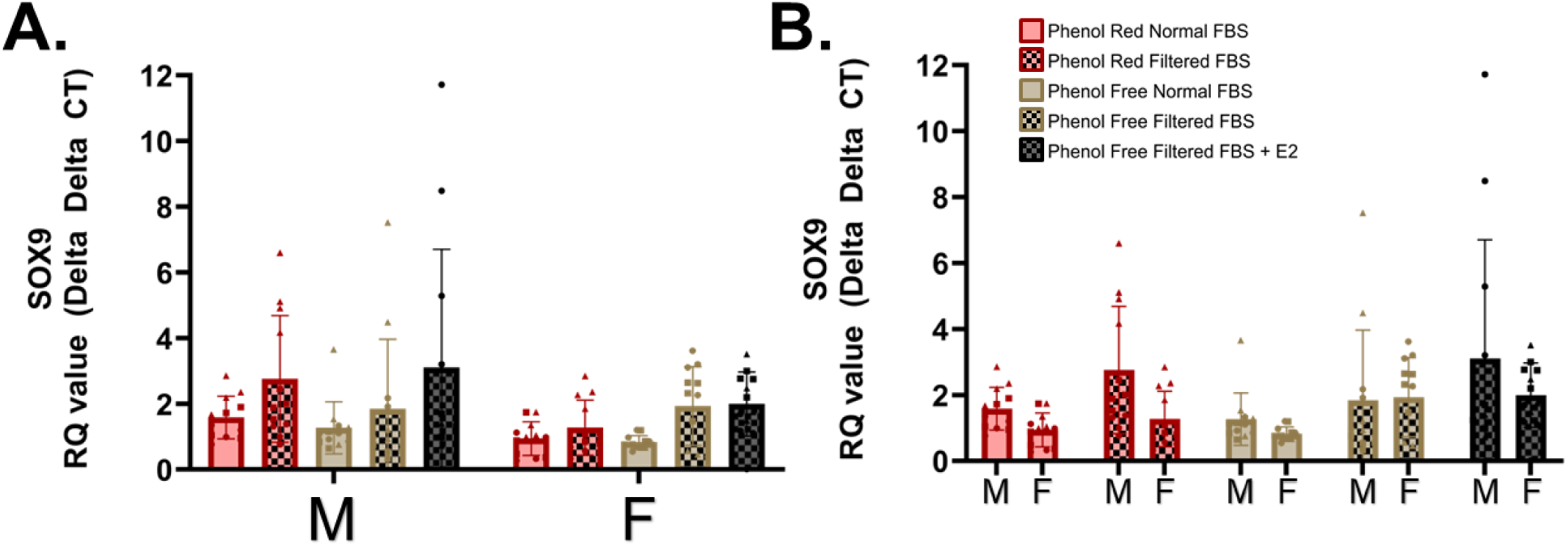
Chondrogenic marker Sox9. Statistical comparisons done within donor sex (A) Statistical comparisons done between donor sexes (B). Analyzed utilizing linear model analysis. SOX9 Linear Model statistics Phenol: P=0.524, FBS: P=0.001, E2: P=0.118, Sex: P=0.015 Interactions: Phenol:FBS: P=0.897, Phenol:Sex: P=0.336, FBS:Sex: P=0.505, E2:Sex: P=0.325, Phenol:FBS:Sex: P=0.296; * P =< .05, ** P =< .01, *** P =< .001, **** P =< .0001. Each dot represents a different well in a tissue culture plate, 4 wells per donor, 3 donors, for a total of 12 points per condition.

## Discussion

### Phenol Red

Phenol red and normal FBS have different, sex specific impacts on male and female cells *in vitro.* In these studies, phenol red is present in the media at a concentration of ∼20 μM compared to the E2 concentration in human serum that ranges from ∼80 pM for males and post-menopausal females (52) to hundreds of nM during pregnancy. Phenol red does bind ERα at a lower Kd (2×10^-5^ M for Phenol Red compared to 2×10^-10^ M for E2) (23) suggesting that phenol red may be acting at an equivalent E2 concentration of 200 pM, being closest to serum estradiol range for menstruating individuals in the follicular or luteal phase (53). Since E2 is known to act in a concentration dependent manner, the relatively high concentration of phenol red may be masking the contributions of E2 in FBS. Phenol red drives proliferation but does not significantly affect metabolism or senescence outputs in this study. Additionally, phenol red does cause female cells to proliferate significantly more than male cells in a manner that is not explained solely by baseline sex differences. Previous results have shown the increase in proliferation due to phenol red requires ERα (23), indicating that phenol red is increasing proliferation through ERα in both male and female cells. Based on the results herein, phenol red is important for proliferation, metabolism, and differentiation and is recommended for use in routine culture. However, because the compound is a non-steroidal estrogen, there are sex-biased effects in most of the metrics assessed and its impact must be considered in applications for stem cell therapy and tissue regeneration.

Phenol red increased osteogenic differentiation as measured by calcium deposition, similar to the results achieved by Lysdahl *et al.* (27). Interestingly, phenol red alone did not have a significant impact on Col10a1 or RUNX2 gene expression. However, a significant increase was observed with normal FBS, similar to the day 14 results from Lysdahl *et al*. In female cells with filtered FBS, phenol red significantly decreased RUNX2 expression compared to the phenol free condition which was also true of its associated osteogenic staining. In female donors, RUNX2 gene expression and calcium staining showed reduced osteogenesis in the phenol red filtered FBS condition and the highest levels of osteogenic differentiation in the phenol free filtered FBS condition, indicating that estrogen deplete media may enhance osteogenesis, a finding also observed in murine MC3T3-E1 osteoblasts (54).

Phenol red also significantly increased adipogenic differentiation capacity. However, in contrast to osteogenic differentiation, this effect was significantly higher in filtered FBS conditions in female donors. This increase from phenol free unfiltered FBS to phenol free filtered FBS was reduced with pM E2 supplementation indicating the E2 component of unfiltered FBS may be the cause of this difference in female donors. Female donors had significantly higher amounts of adipogenic differentiation compared to male donors in phenol red conditions. The increased adipogenic staining was accompanied by an increase in FABP4 expression in filtered FBS conditions. While this is the first data illustrating the impact of phenol red on adipogenic differentiation, previous studies have looked at the impact of E2 on adipogenic differentiation, largely finding a linear increase in adipogenic staining with increasing E2 treatment (55). Both of these results run counter to Okazaki et al. 2002, however those studies were performed in murine bone marrow derived-MSCs (56). Overall, phenol red significantly alters hMSCs, albeit not detrimentally, through an increase in proliferation with minimal impact on senescence staining or other markers of cellular health, indicating its routine inclusion in studies where E2 signaling is not of a particular concern may be desirable.

### E2 in FBS

Filtered FBS had the greatest impact on hMSC metabolism, senescence, and differentiation. Specifically, the reductive capacity of cells was significantly reduced in filtered FBS. When analyzed via Seahorse XF analyzer, the basal metabolic rate of hMSCs varied drastically in female, but not male cells. The metabolic profiles of hMSCs can be indicative of their stem-like potential (57). Specifically, hMSCs with a higher reliance on glycolytic metabolism have been shown to exhibit enhanced stem-like properties (58). On the other hand, hMSCs with a higher reliance on oxidative phosphorylation may signal lineage specification indicating that normal FBS maybe be causing differentiation in hMSCs prior to the intentional induction of differentiation cues, particularly in female cells. This consideration is critical, as committing to lineage differentiation prior to the intention of the researcher may cause issues for lineage specific work, obfuscate differentiation timelines, and further prevent hMSCs from acting in their trophic capacities to reduce inflammation. Male cells did not exhibit significant metabolic changes in their basal metabolic rates in any of the media formulations. Of note, individual female donors were highly variable in this data, with some donors having extremely low OCR/ECAR even in the phenol red normal FBS condition. These patterns may be altered in donors of older cells, as other research has indicated that young female cells do maintain a lower OCR/ECAR ratio than male cells, but that can change as they are aged and stressed (59). Filtered FBS rescued the female donors to a more stem-like metabolic profile, with the donor BM04 also showing a reduction in senescence in the phenol red, filtered FBS condition in comparison to the phenol red, unfiltered FBS condition (**Supplemental Figure 5**). Male cells also showed a reduction in senescence-associated β-galactosidase staining in the phenol red, but not the phenol free condition in response to filtered FBS.

Regarding osteogenic differentiation of male and female hMSCs, unfiltered FBS resulted in an increase in alizarin red staining compared to filtered FBS in phenol red conditions. Additionally, only in unfiltered FBS conditions did male cells see a significant increase in Col10a1 expression, matching the significant differences seen in osteogenic staining. However, this effect was largely driven by a single donor. FBS filtration did not significantly impact RUNX2 expression. Phenol free media did not have a significant impact on osteogenic differentiation, however, phenol red as a parameter did significantly contribute to the response of these cells to the FBS conditions. These results are contrary to Lysdahl *et al.* (60) which showed that osteogenic differentiation in phenol red free conditions significantly increased – albeit, only in a single male donor. Other previous work performed in chondrocytes demonstrated that unfiltered FBS was needed for E2 treatment to impact osteogenic differentiation. These results agree with our data where unfiltered FBS conditions had significant increases in expression of osteogenic markers (61). Our data shows that phenol red overall increases Col10a1 expression as well as osteogenic staining at 2 weeks. Similarly, Lysdahl et al. showed increased osteogenesis on day 14 with phenol red, that then reversed at 21 days, indicating these results may shift in longer-term culture.

Increases in lipid droplet accumulation were only observed in phenol free conditions with female donors with the adipogenic differentiation media. All other conditions showed no significant differences between filtered and unfiltered FBS in matched phenol conditions. Filtered FBS, particularly in phenol red conditions, resulted in higher adipogenic staining and expression of FABP4 for female cells compared to the phenol free normal FBS and E2 control conditions. Additionally female donors with filtered FBS experienced significantly higher baseline staining in phenol red and phenol red free conditions. Unfiltered FBS decreases the ‘stem like metabolic profile’ of female cells, is the only driver of senescence in both male and female cells and increases osteogenic differentiation. As such, unfiltered FBS should only be utilized where rapid osteogenesis is the primary goal, in all other metrics filtered FBS exhibits superior stemness and overall cellular health.

### Sex Differences

Sex differences are summarized in **Figure 12**. To properly assess differences due to sex, we must consider both the inherent differences due to sex and the contribution of sex to the differential response of cells to the culture conditions via interaction parameters using linear model assessment. Linear model parameters (**Table 2**) for proliferation indicate that there are both inherent differences in proliferation within the same treatment groups, and that the treatment combined with sex create a significantly different effect than what the sex differences or treatment alone would cause. Metabolic outcomes exhibited baseline differences by donor sex, however donor sex did not significantly alter metabolic response to any other parameter. Senescence associated staining was not altered by donor sex in this study, but was rather dependent on the individual donor. Direct differentiation of donor cells showed that adipogenic differentiation has both baseline sex differences, and a significant contribution from sex to the reaction to exogenous estrogens. Previous research has detailed increased hMSC adipogenic differentiation with E2 treatment *in vitro*. However, there is *in vivo* evidence that in older individuals, E2 may suppress adipogenic differentiation (55,62), highlighting the importance of considering both age and E2 concentration. Osteogenic differentiation showed a significant baseline difference in sex and significant interactions between FBS and donor sex, as well as phenol red, FBS, and donor sex. SOX9, Col10a1, and FABP4 expression were all significantly different based on donor sex for baseline expression, but not significantly different in response to treatment. RUNX2 expression, which mapped well onto the osteogenic staining results, indicating a significant difference in both baseline sex differences and response to exogenous estrogens based on donor sex. Given the RUNX2 and alizarin red data alignment, particularly for female cells, it is fair to conclude that osteogenic differentiation is significantly affected by a sex specific response to exogenous estrogens.

**Figure 12.**
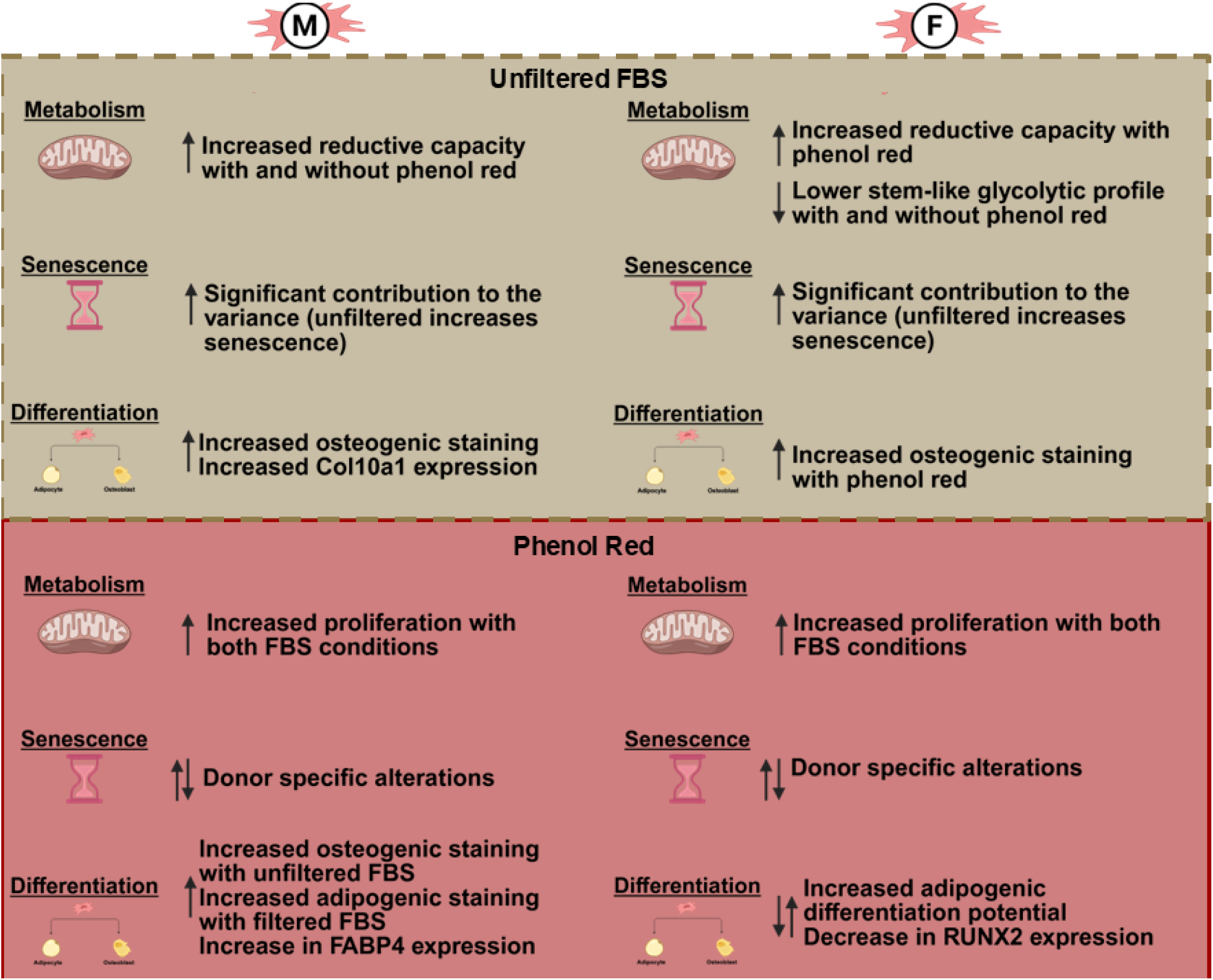
Visual summary of observed differences by sex based on presence of phenol red and unfiltered FBS in culture media.

These results may be partially explained by E2 induced epigenetic regulation as it has been shown to impact osteogenic and adipogenic transcription factors, although this is muddied by the increase in methylation in ‘aged’ hMSCs as defined by increased β-galactosidase staining, which as shown here is also altered by unfiltered FBS (63). Other effects of E2 may similarly be contradictory, where lncRNA-H19 is known to downregulate proliferation and osteogenesis in hMSCs, but only when E2 is present (64). However, E2 alone has also been known to increase osteogenesis (55).

Overall, there are many differences based on hMSCs exposure to estrogens in the media, with many of the outcomes having baseline sex differences and differential response based on sex. In hMSC applications where the desired culture outcomes are typically quick proliferation, low levels of senescence, and limited differentiation, our results show that phenol red media with charcoal dextran filtered FBS is optimal. However, phenol red may obfuscate the ability to interrogate differences in estrogen signaling and should be excluded from studies investigating hormone signaling. Additionally, phenol red may increase senescence associated staining in individual donors, so pre-screening hMSCs for this response may be prudent. Clinically, it may be advantageous to culture hMSCs in media with filtered FBS, as this will minimize metabolic shifts and maximize the stem cell like phenotype of the cells.

While this study represents a comprehensive view of the unintentional biases created in hMSC culture when using traditional culture media with estrogens, there are some critical limitations. Charcoal dextran filtering is common for the removal of hormones such as E2, however it is not specific for E2 and as such there may be other hormones, such as triiodothyronine (T3), or other compounds that were filtered out that may have a dominant impact. Because of this knowledge, we re-supplemented specifically with pM levels of E2 to match the reported serum levels, to determine whether observations with CD-FBS were E2 specific. Also, culture past 14 days to assess lineage specific differentiation may provide more information about the temporal action of these compounds presently found in cell culture media. Finally, the observed effects are primarily applicable to young hMSC donors (<23 years), and it’s recognized that aged donors may have marked changes in their overall cellular health and differentiation potential.

## Conclusion

Human MSCs are a promising cell source for regenerative cell therapy. However, much work needs to be done to understand how sex differences impact their basic cell processes, especially in relation to regeneration. One of the variables that needs to be well characterized is the differential response of cells to hormones, specifically estrogens. To understand the differential response of cells to E2, the baseline impact of exogenous E2 and E2-mimetic compounds commonly present in cell culture on hMSC behavior must be understood. In this study, we demonstrated that phenol red significantly increases proliferation with minimal deleterious effects on cells, however any epigenetic effects are still unknown and warrant future study. Unfiltered fetal bovine serum reduces metabolic stemness and increases senescence-associated staining while also altering osteogenic and chondrogenic gene expression. Donor sex significantly impacted every aspect of cellular health and phenotype tested, except for PPARG expression and senescence. Sex specific responses beyond differences based on sex to exogenous forms of E2 were noted in proliferation, osteogenic and adipogenic differentiation capacity, and COL1a1, COL10a1 and RUNX2 expression. Future work should be done to investigate the long-term epigenetic effects of exogenous estrogens as E2 is known to affect DNA methylation and other epigenetic regulatory mechanisms that may be altered long-term in culture based on limited exposure. Moving forward, these results should be considered whenever designing a study investigating sex-biased effects.

## Supporting information

Supplemental Graphs Code Text and Images

## Acknowledgments

We would like to acknowledge funding from NIH NIGMS Maximizing Investigators Research Award (R35GM143081) and the University of Washington Institute for Stem Cell and Regenerative Medicine (ISCRM) startup funds. The content is solely the responsibility of the authors and does not necessarily represent the official views of the funders including the National Institutes of Health. We would like to acknowledge the Diabetes Research Center with NIH P30DK017047 funding for their assistance in making the Seahorse XF Analyzer available for usage and training on the instrumentation. We would also like to thank members of the Robinson Lab for their commentary on data and aiding in editing this paper including Susana P. Simmonds Bohorquez, Katherine L. Meinhold, Kyley Burkey, and Janie Johnson and Dr. Ron Kwon for his assistance in statistical methods. The University of Washington acknowledges the Coast Salish peoples of this land, the land which touches the shared waters of all tribes and bands within the Suquamish, Tulalip and Muckleshoot nations.

Corresponding Author: Jennifer L Robinson

