## Supplemental Graphs Code Text and Images for "Media Matters: Phenol Red and Fetal Bovine Serum Estrogen in Traditional Cell Culture Media Influence Human Mesenchymal Stromal Cell (hMSC) Processes and Differentiation in a Sex-Biased Manner"

Supplementary Figure 1: Legend

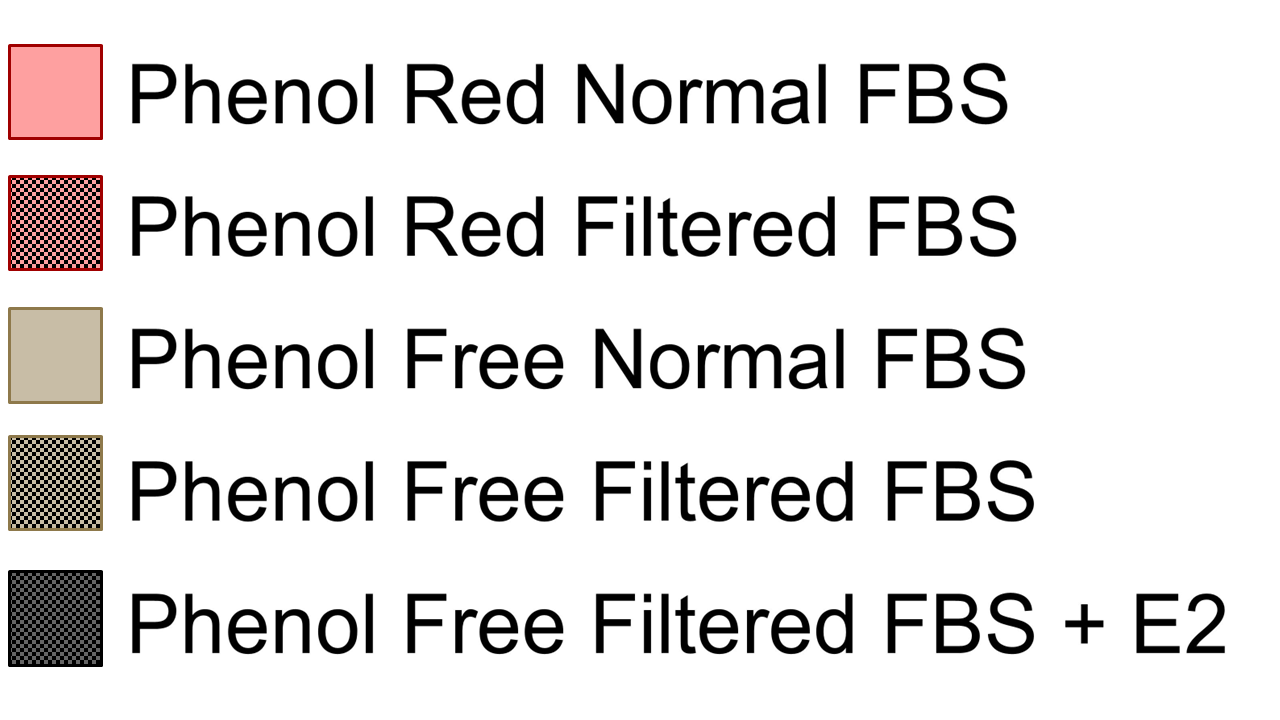
**Figure 1**: Color and shading legend for all supplementary figures

Supplementary Figure 2: Proliferation

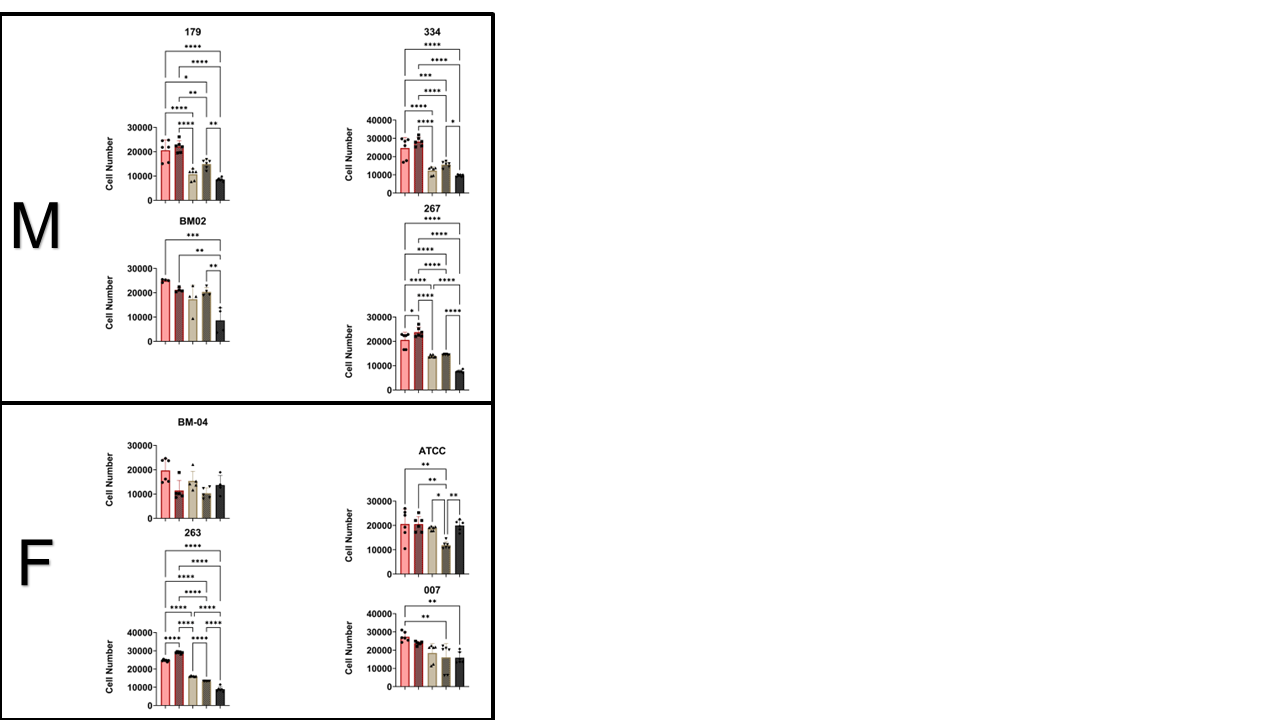

**Figure 2**: Individual Graphs for Proliferation assayed VIA Picogreen Statistical comparisons done within individual donors. All data tested for normality utilizing the skewness test (s < .5) and analyzed utilizing 2-way ANOVA, and Bonferroni multiple comparison testing. ANOVA Statistics * P =< .05, ** P =< .01, *** P =< .001, **** P =< .0001. Each dot represents a different well in a tissue culture plate, 3 wells per donor, 4 donors, for a total of 12 points per condition.

Supplementary Figure 3: Metabolism by Prestoblue

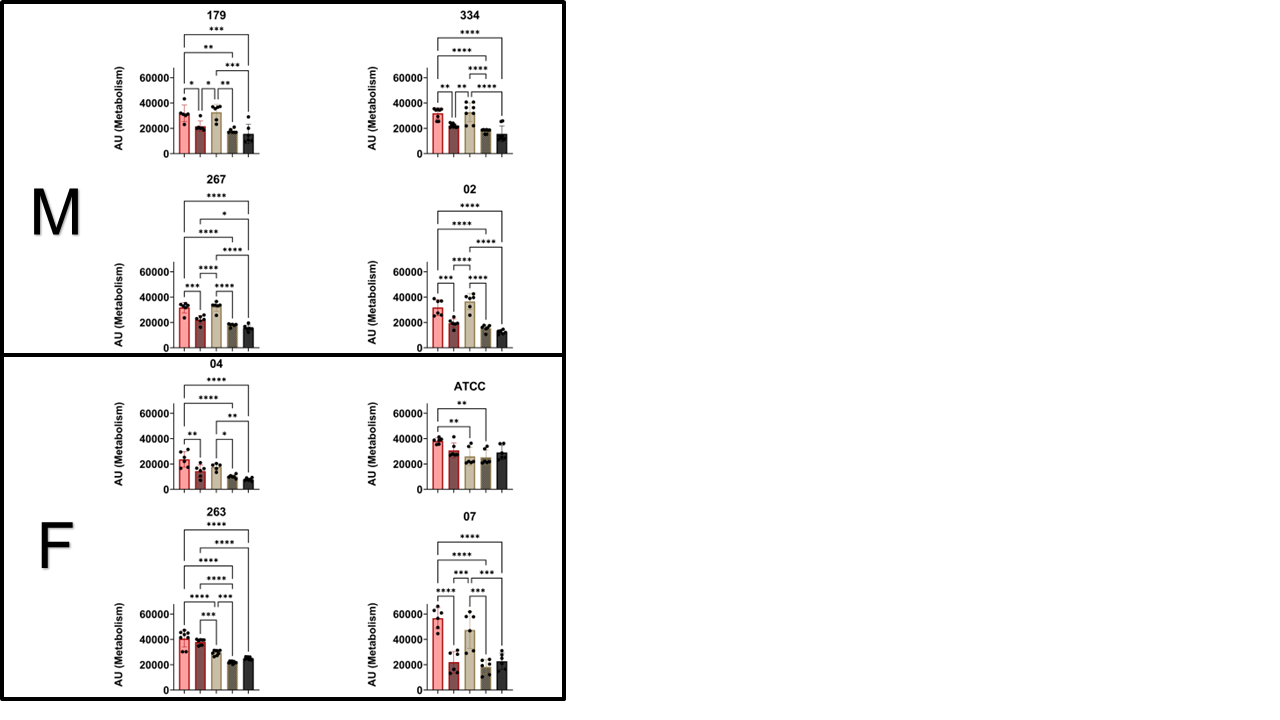

**Figure 3**: Individual Graphs for Metabolism assayed VIA Prestoblue Statistical comparisons done within individual donors. All data tested for normality utilizing the skewness test (s < .5) and analyzed utilizing 2-way ANOVA, and Bonferroni multiple comparison testing. ANOVA Statistics * P =< .05, ** P =< .01, *** P =< .001, **** P =< .0001. Each dot represents a different well in a tissue culture plate, 6-8 wells per donor, 4 donors per sex.

Supplementary Figure 4: Seahorse Basal Metabolic Rate

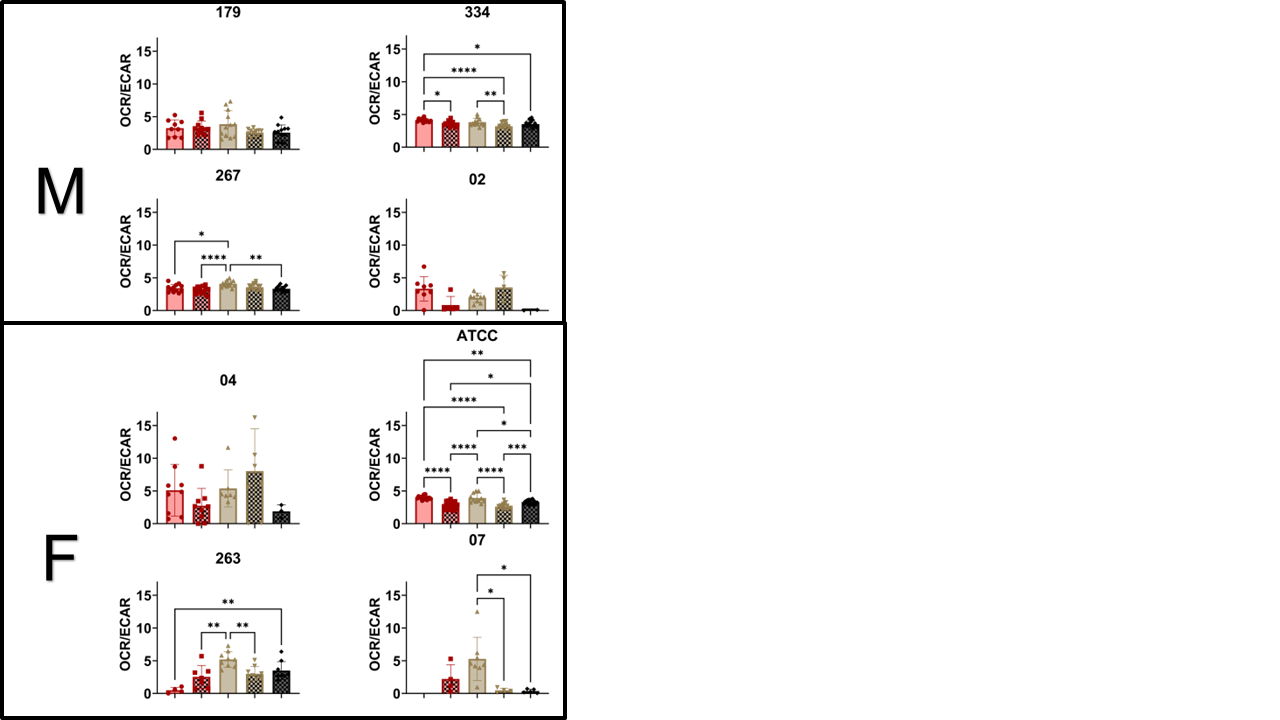

**Figure 4**: Individual Graphs for Metabolism assayed VIA Seahorse Basal Metabolic Rate Statistical comparisons done within individual donors. All data tested for normality utilizing the skewness test (s < .5) and analyzed utilizing 2-way ANOVA, and Bonferroni multiple comparison testing. ANOVA Statistics * P =< .05, ** P =< .01, *** P =< .001, **** P =< .0001. Each dot represents a different well in a tissue culture plate, 6-12 wells per donor, 4 donors per sex.

Supplementary Figure 5: β-Galactosidase Senescence Associated Staining

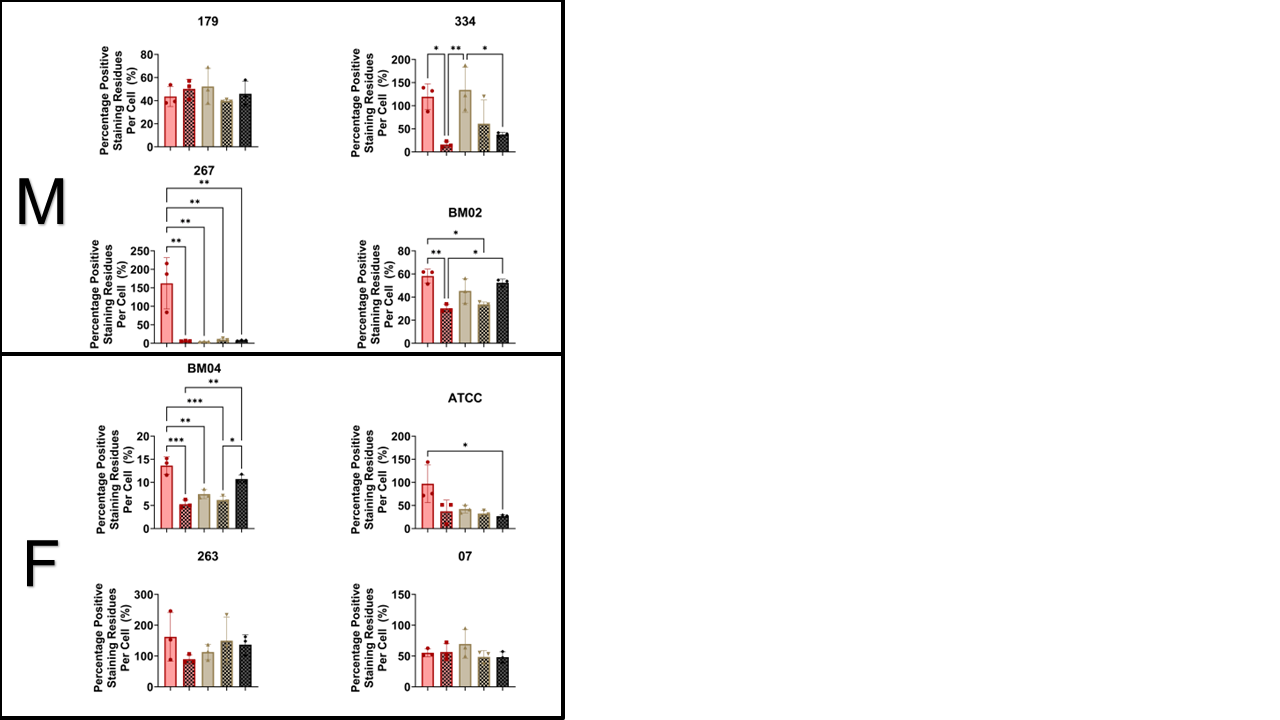

**Figure 5**: Individual Graphs for Senescence assayed β-Galactosidase Staining. Statistical comparisons done within individual donors. All data tested for normality utilizing the skewness test (s < .5) and analyzed utilizing 2-way ANOVA, and Bonferroni multiple comparison testing. ANOVA Statistics * P =< .05, ** P =< .01, *** P =< .001, **** P =< .0001. Each dot represents a different well in a tissue culture plate, 3 wells per donor, 4 donors per sex.

Supplementary Figure 6: Alizarin Red Stain

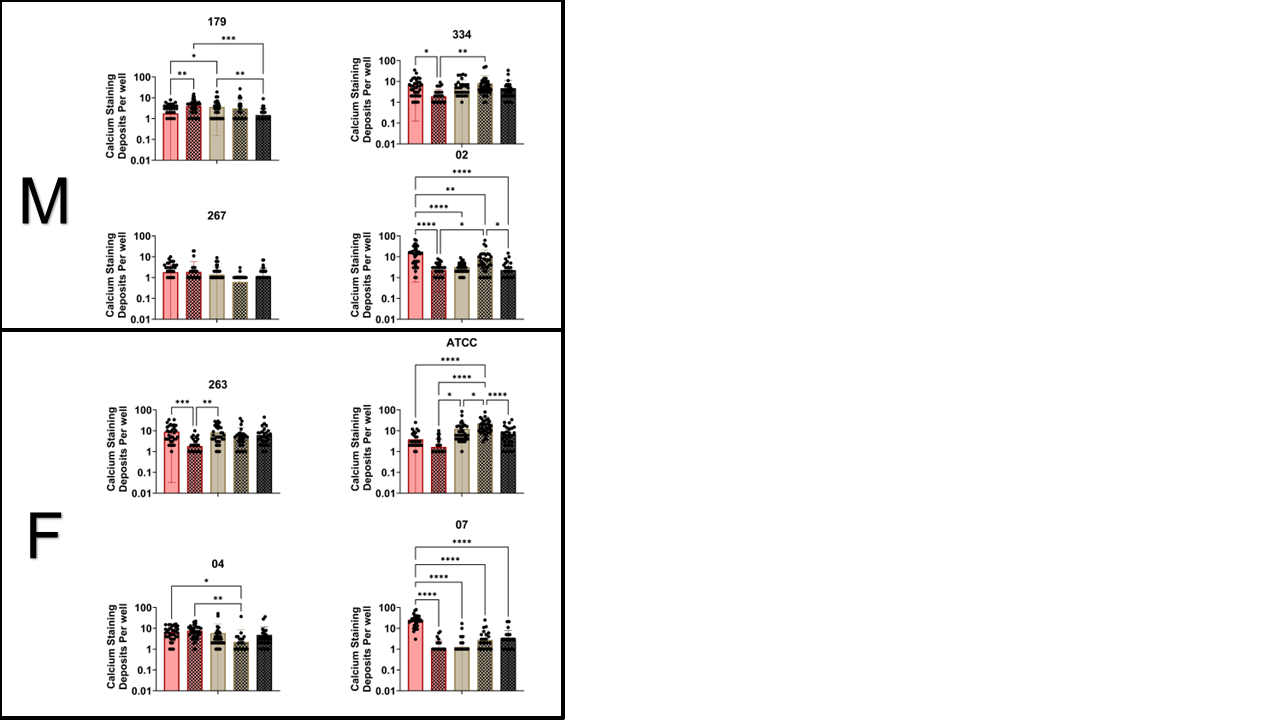

**Figure 6**: Individual Graphs for Osteogenic Differentiation assayed Alizarin Red Staining. Statistical comparisons done within individual donors. All data tested for normality utilizing the skewness test (s < .5) and analyzed utilizing 2-way ANOVA, and Bonferroni multiple comparison testing. ANOVA Statistics * P =< .05, ** P =< .01, *** P =< .001, **** P =< .0001. Each dot represents a different image from a well in a tissue culture plate, 36-50 images per well per donor, 4 donors per sex.

Supplementary Figure 7: Oil Red O Stainin

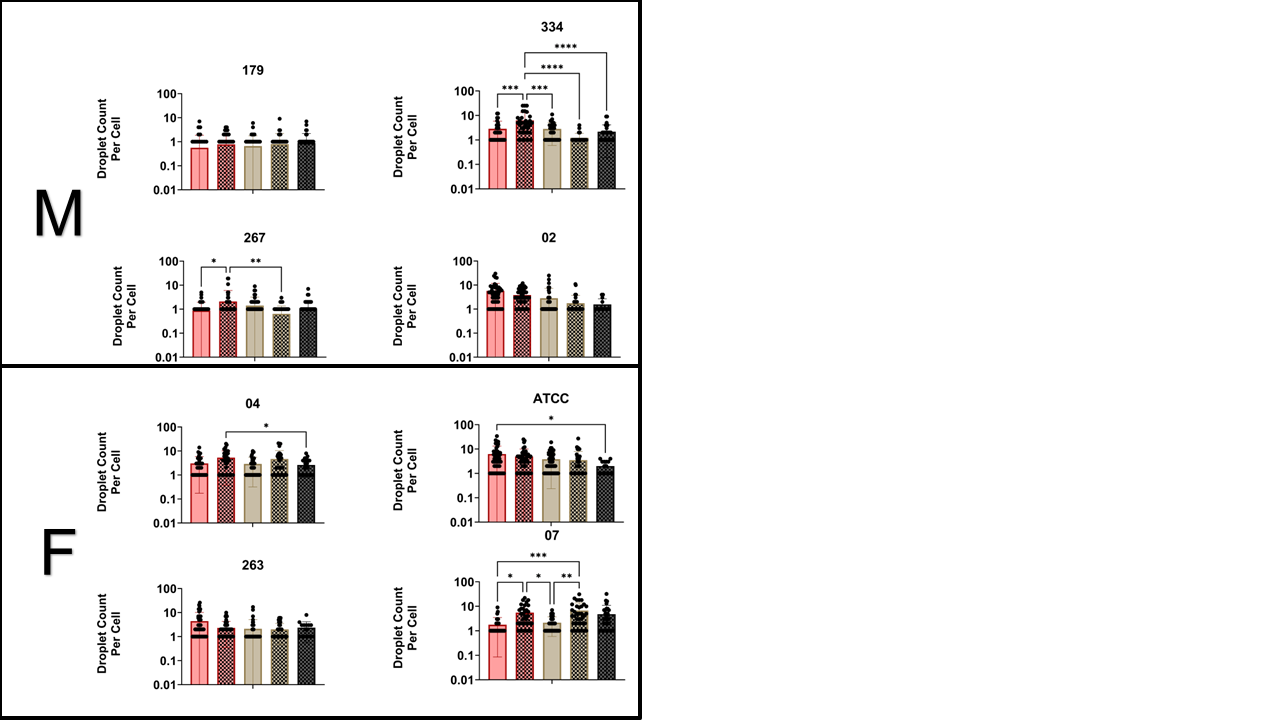

**Figure 7**: Individual Graphs for Adipogenic Differentiation assayed Oil Red O Staining. Statistical comparisons done within individual donors. All data tested for normality utilizing the skewness test (s < .5) and analyzed utilizing 2-way ANOVA, and Bonferroni multiple comparison testing. ANOVA Statistics * P =< .05, ** P =< .01, *** P =< .001, **** P =< .0001. Each dot represents a different image from a well in a tissue culture plate, 36-50 images per well per donor, 4 donors per sex.

Supplementary Figure 8: ESR1 qPCR

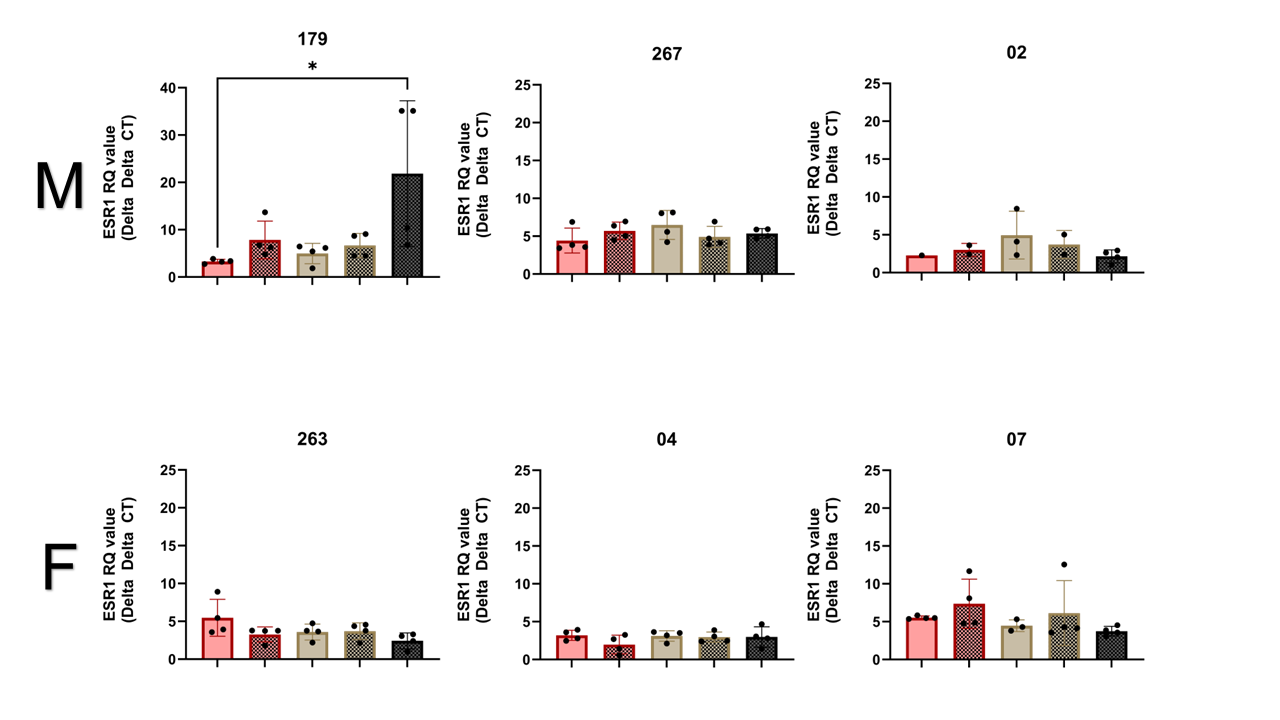
**Figure 8**: Individual Graphs for ESR1 expression assayed qPCR. Statistical comparisons done within individual donors. All data tested for normality utilizing the skewness test (s < .5) and analyzed utilizing 2-way ANOVA, and Bonferroni multiple comparison testing. ANOVA Statistics * P =< .05, ** P =< .01, *** P =< .001, **** P =< .0001. Each dot represents a different image from a well in a tissue culture plate, 4 wells per donor, 4 donors per sex.

Supplementary Figure 9: COL1a1 qPCR

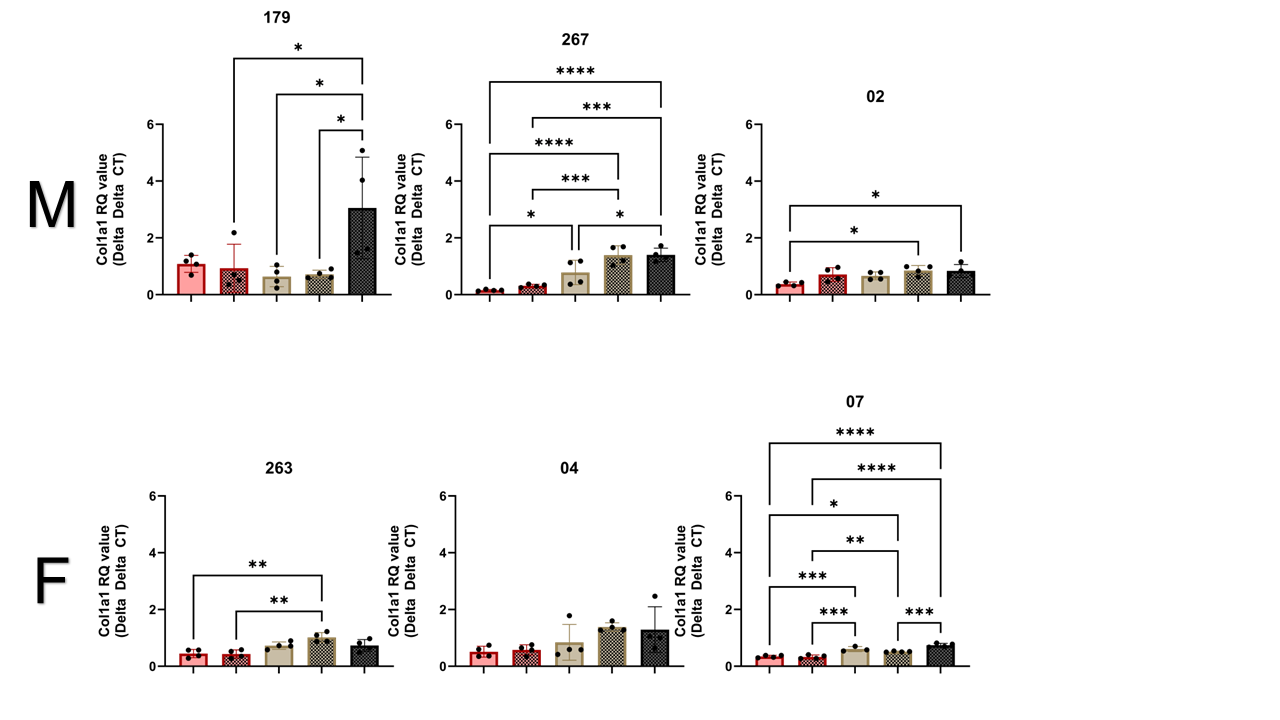
**Figure 9**: Individual Graphs for COL1a1 expression assayed qPCR. Statistical comparisons done within individual donors. All data tested for normality utilizing the skewness test (s < .5) and analyzed utilizing 2-way ANOVA, and Bonferroni multiple comparison testing. ANOVA Statistics * P =< .05, ** P =< .01, *** P =< .001, **** P =< .0001. Each dot represents a different image from a well in a tissue culture plate, 4 wells per donor, 4 donors per sex.

Supplementary Figure 10: RUNX2 qPCR

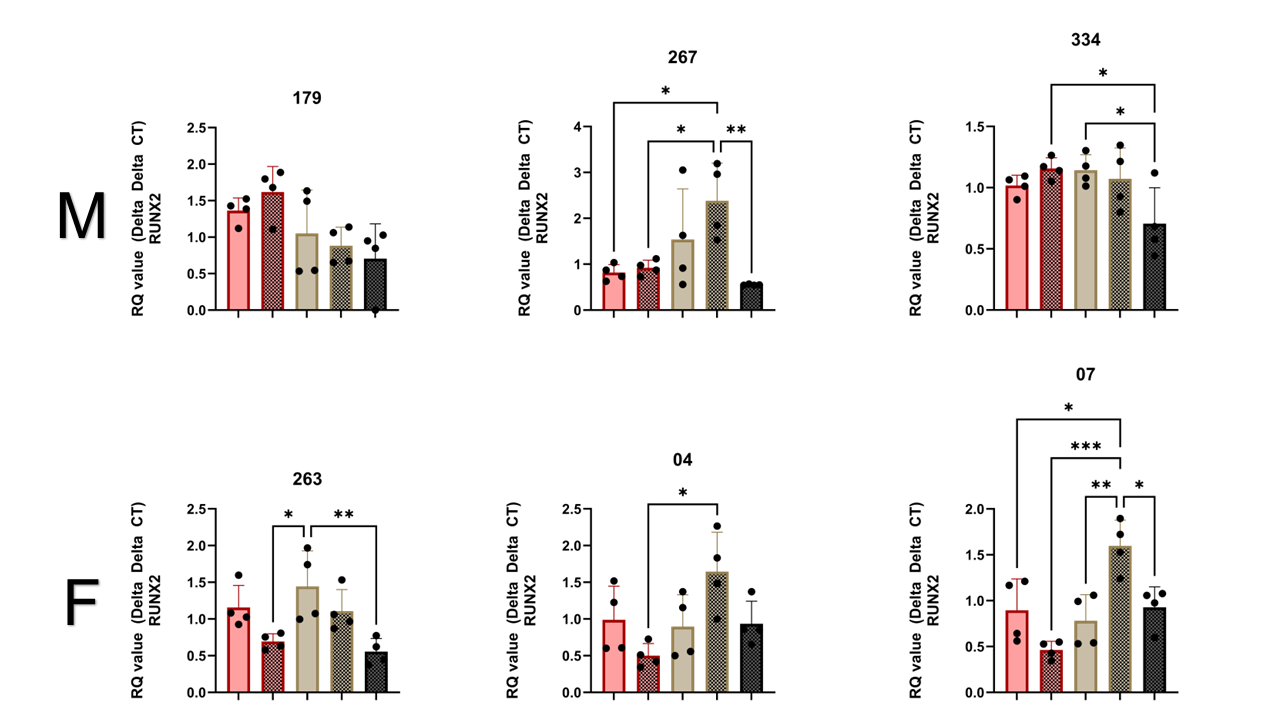
**Figure 10**: Individual Graphs for RUNX2 expression assayed qPCR. Statistical comparisons done within individual donors. All data tested for normality utilizing the skewness test (s < .5) and analyzed utilizing 2-way ANOVA, and Bonferroni multiple comparison testing. ANOVA Statistics * P =< .05, ** P =< .01, *** P =< .001, **** P =< .0001. Each dot represents a different image from a well in a tissue culture plate, 4 wells per donor, 4 donors per sex.

Supplementary Figure 11: COL10a1 qPCR

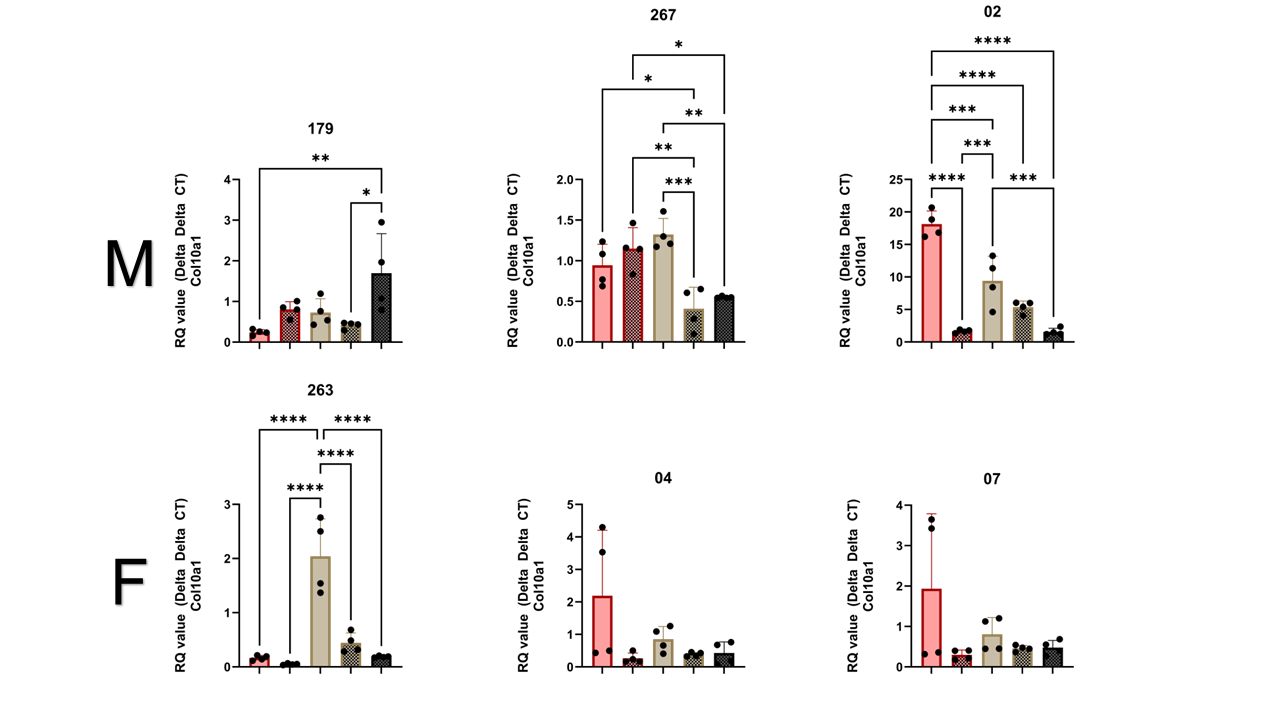
**Figure 11**: Individual Graphs for COL10a1 expression assayed qPCR. Statistical comparisons done within individual donors. All data tested for normality utilizing the skewness test (s < .5) and analyzed utilizing 2-way ANOVA, and Bonferroni multiple comparison testing. ANOVA Statistics * P =< .05, ** P =< .01, *** P =< .001, **** P =< .0001. Each dot represents a different image from a well in a tissue culture plate, 4 wells per donor, 4 donors per sex.

Supplementary Figure 12: FABP4 qPCR

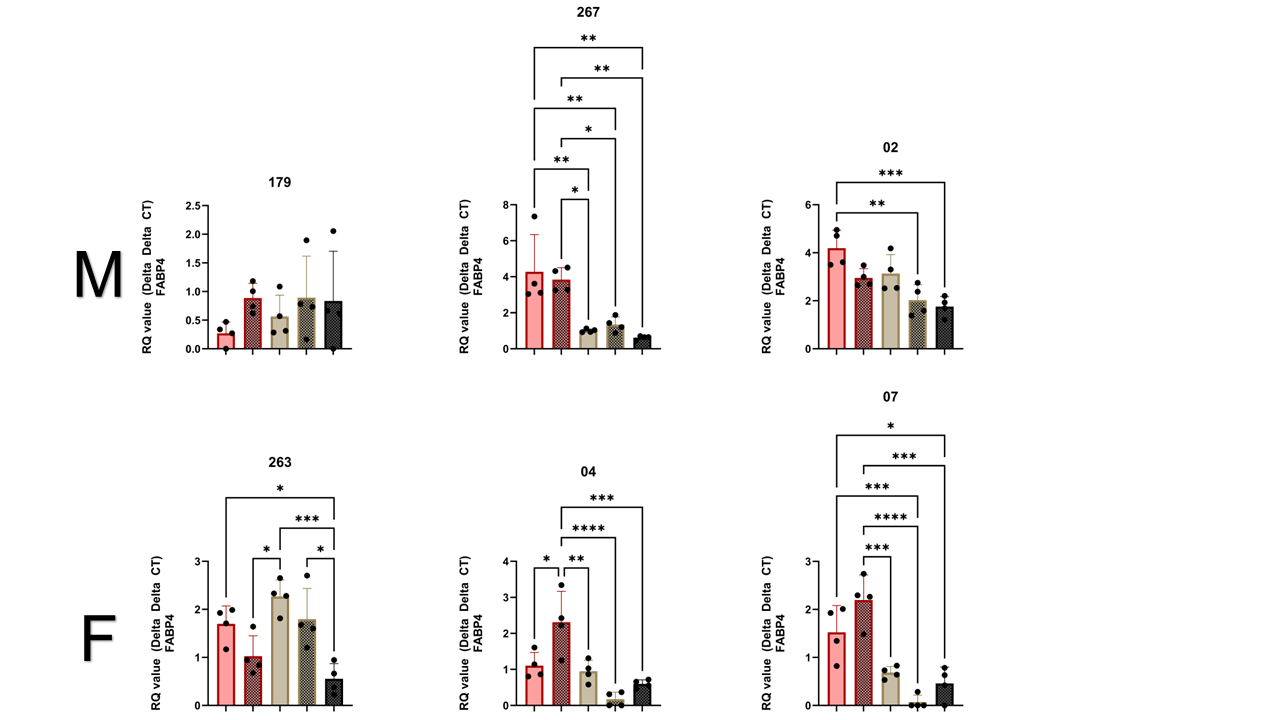
**Figure 12**: Individual Graphs for FABP4 expression assayed qPCR. Statistical comparisons done within individual donors. All data tested for normality utilizing the skewness test (s < .5) and analyzed utilizing 2-way ANOVA, and Bonferroni multiple comparison testing. ANOVA Statistics * P =< .05, ** P =< .01, *** P =< .001, **** P =< .0001. Each dot represents a different image from a well in a tissue culture plate, 4 wells per donor, 4 donors per sex.

Supplementary Figure 13: PPARγ qPCR

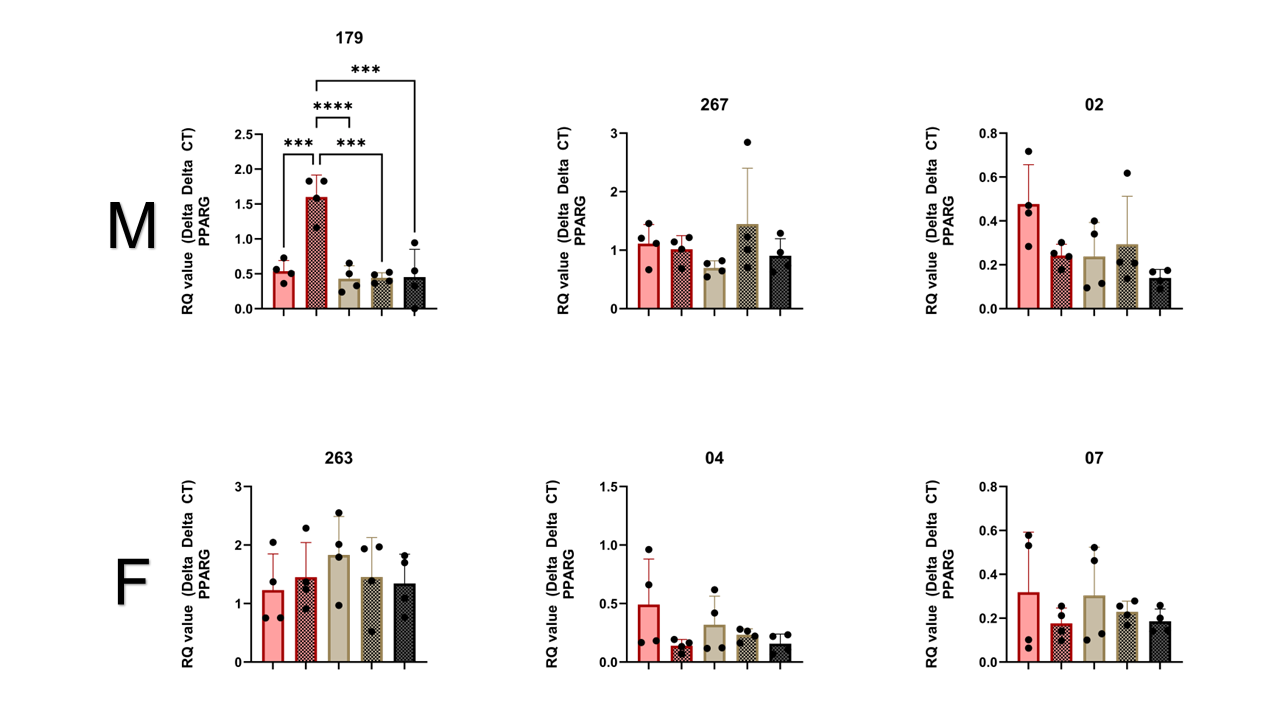
**Figure 13**: Individual Graphs for PPARγ expression assayed qPCR. Statistical comparisons done within individual donors. All data tested for normality utilizing the skewness test (s < .5) and analyzed utilizing 2-way ANOVA, and Bonferroni multiple comparison testing. ANOVA Statistics * P =< .05, ** P =< .01, *** P =< .001, **** P =< .0001. Each dot represents a different image from a well in a tissue culture plate, 4 wells per donor, 4 donors per sex.

Supplementary Figure 14: SOX9 qPCR

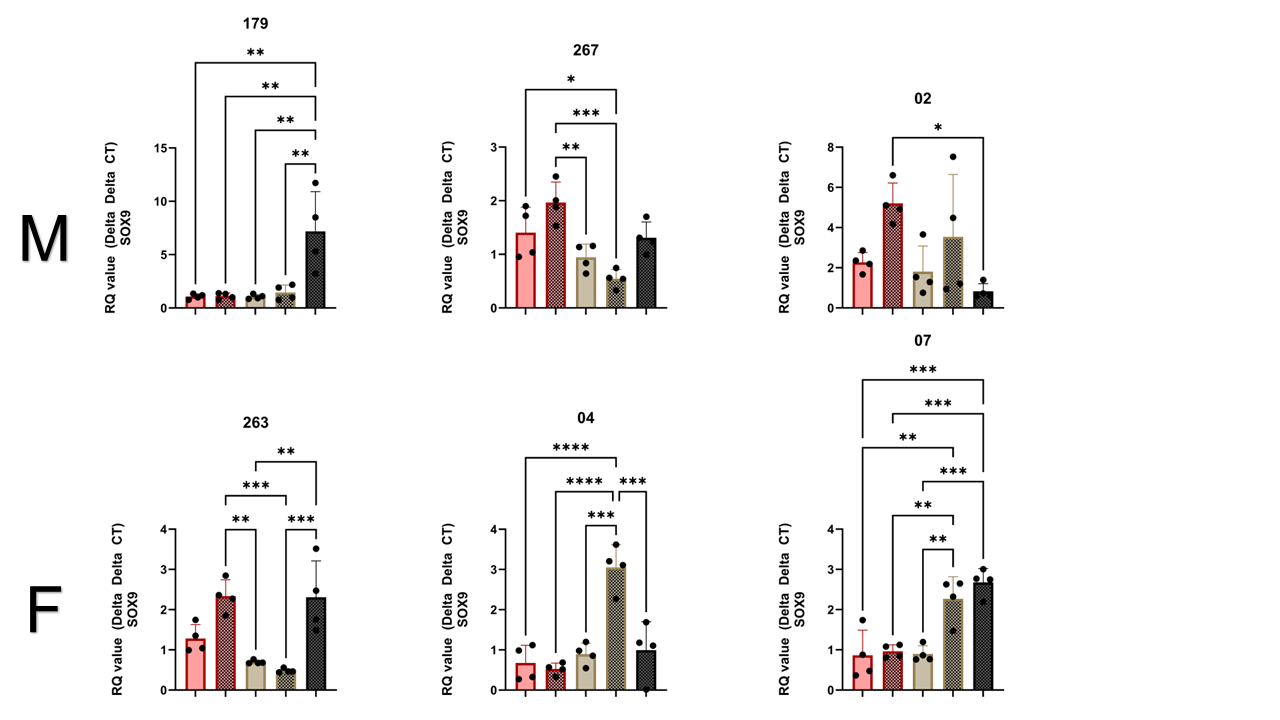
**Figure 14**: Individual Graphs for SOX9 expression assayed qPCR. Statistical comparisons done within individual donors. All data tested for normality utilizing the skewness test (s < .5) and analyzed utilizing 2-way ANOVA, and Bonferroni multiple comparison testing. ANOVA Statistics * P =< .05, ** P =< .01, *** P =< .001, **** P =< .0001. Each dot represents a different image from a well in a tissue culture plate, 4 wells per donor, 4 donors per sex.

Supplementary Text 1: Macro Code for IJM using FIJI Senescence

i = 1;

while (i <= 4063) {

//4063

sample = "Sample (" + i + ").jpg";

path = "path to file"+sample;

open(path);

run("Gaussian Blur...", "sigma=5");

run("Colour Deconvolution", "vectors=RGB");

close;

close;

run("Auto Threshold", "method=Triangle");

run("8-bit");

sample_Colour_1 = "Sample (" + i + ").jpg-(Colour_1)";

imageCalculator("Add create", sample, sample_Colour_1);

run("Colour Deconvolution", "vectors=[User values] [r1]=124 [g1]=155 [b1]=111 [r2]=152 [g2]=0.00000 [b2]=255 [r3]=128 [g3]=137 [b3]=255");

close();

run("Gaussian Blur...", "sigma=5");

run("Auto Local Threshold", "method=Phansalkar radius=500 parameter_1=0 parameter_2=0");

run("Analyze Particles...", "size=2000-Infinity circularity=0.20-1.00 show=Overlay display clear summarize");

run("Close All");

i++;

}

Supplementary Text 2: Macro Code for IJM using FIJI Alizarin Red

i = 1;

while (i <= 1650) {

sample = "Sample (" + i + ").jpg";

path = "path to file" + "Sample (" + i + ").jpg";

open(path);

run("Enhance Contrast", "saturated=0.35");

setMinAndMax(0, 255);

run("Gaussian Blur...", "sigma = 2");

run("Colour Deconvolution", "vectors=[User values] [r1]=0.48696724 [g1]=0.57941777 [b1]=0.6535579 [r2]=0.0347683 [g2]=0.5229714 [b2]=0.8516408 [r3]=0.13686699 [g3]=0.39844745 [b3]=0.90692174");

close();

run("Auto Threshold", "method=RenyiEntropy");

run("Invert");

run("Analyze Particles...", " circularity=0.50-1.00 display clear summarize");

run("Close All");

i++;

}

Supplementary Text 3: Macro Code for IJM using FIJI Oil Red O

i = 1;

while (i <= 1650) {

sample = "Sample (" + i + ").jpg";

path = "path to file” + "Sample (" + i + ").jpg";

open(path);

run("Gaussian Blur...", "sigma=5");

run("Colour Deconvolution", "vectors=[User values] [r1]=0.48696724 [g1]=0.57941777 [b1]=0.6535579 [r2]=0.0347683 [g2]=0.5229714 [b2]=0.8516408 [r3]=0.13686699 [g3]=0.39844745 [b3]=0.90692174");

close();

run("Gaussian Blur...", "sigma=5");

run("Auto Threshold", "method=RenyiEntropy");

run("Invert");

run("Analyze Particles...", "size=100-Infinity circularity=0.30-1.00 show=Overlay display clear summarize");

run("Close All");

i++;

}

Table 1: Review of Clinical Trials and their Inclusion of Exogenous Forms of Estrogen

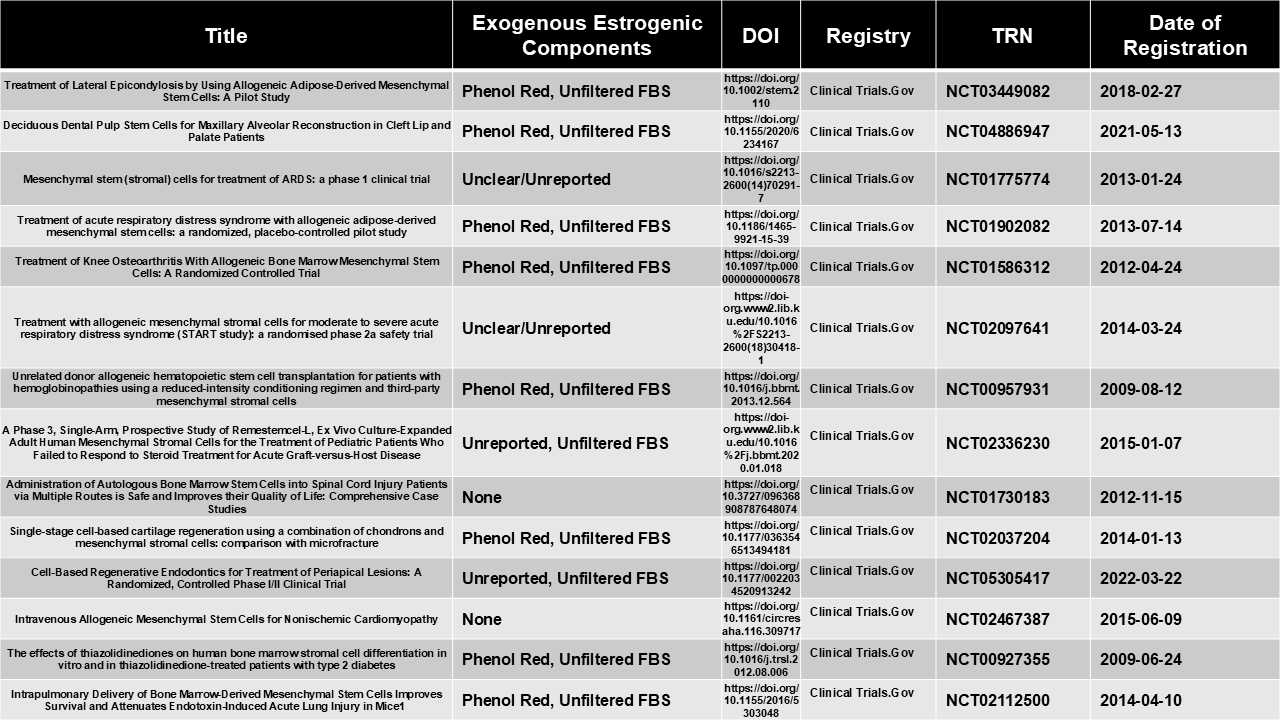

Image 1: COA of Hormone Content in Unfiltered and Filtered FBS

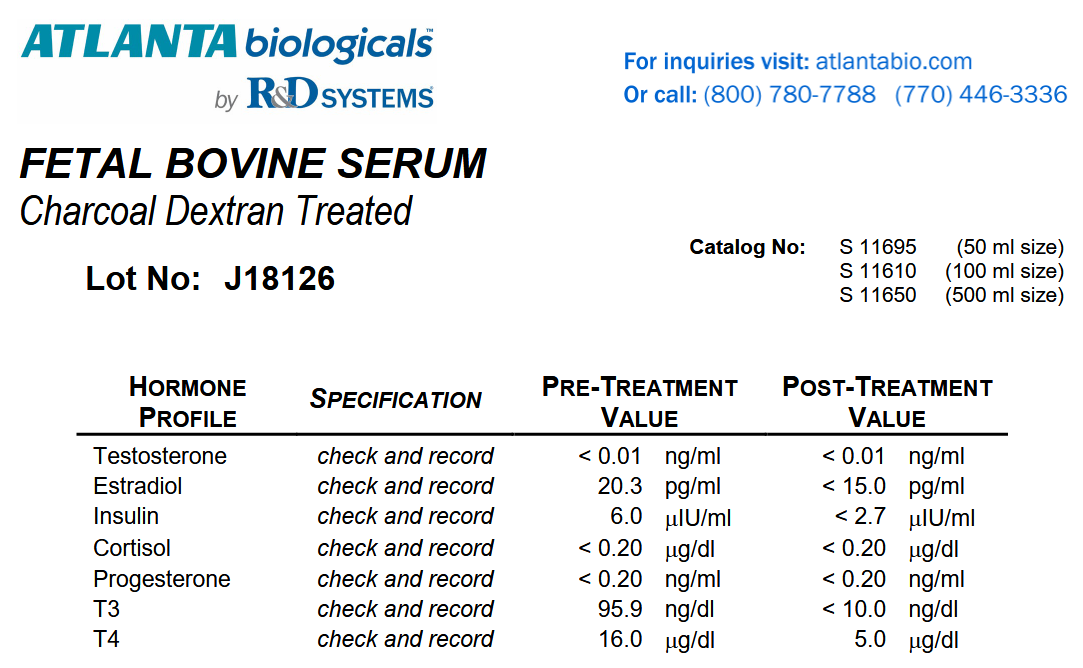
